# Function and firing of the *Streptomyces coelicolor* contractile injection system requires the membrane protein CisA

**DOI:** 10.1101/2024.06.25.600559

**Authors:** Bastien Casu, Joseph W. Sallmen, Peter E. Haas, Govind Chandra, Pavel Afanasyev, Jingwei Xu, Martin Pilhofer, Susan Schlimpert

**Affiliations:** Department of Biology, Institute of Molecular Biology & Biophysics, Eidgenössische Technische Hochschule Zürich, Otto-Stern-Weg 5, 8093 Zürich, Switzerland; John Innes Center, Department of Molecular Microbiology, Norwich Research Park, Norwich, NR4 7UH, United Kingdom; Cryo-EM Knowledge Hub, Eidgenössische Technische Hochschule Zürich, Otto-Stern-Weg 3, 8093 Zürich, Switzerland; Centre for Microbial Interactions, Norwich Research Park, Norwich, NR4 7UH, United Kingdom

**Keywords:** bacterial development/cellular microbiology/contractile phage tail-like structures/regulated cell death/multicellularity

## Abstract

Bacterial contractile injection systems (CIS) are phage tail-like macromolecular complexes that mediate cell-cell interactions by injecting effector proteins into target cells. CIS from *Streptomyces coelicolor* (CIS^Sc^) are localized in the cytoplasm. Under stress, they induce cell death and impact the *Streptomyces* life cycle. It remains unknown, however, whether CIS^Sc^ require accessory proteins to directly interact with the cytoplasmic membrane to function.

Here, we characterize the putative membrane adaptor CisA, a conserved factor in *CIS* gene clusters across *Streptomyces* species. We show by cryo-electron tomography imaging and *in vivo* assays that CIS^Sc^ contraction and function depend on CisA. Using single-particle cryo-electron microscopy, we provide an atomic model of the extended CIS^Sc^ apparatus; however, CisA is not part of the complex. Instead, our findings show that CisA is a membrane protein with a cytoplasmic N-terminus predicted to interact with CIS^Sc^ components, thereby providing a possible mechanism for mediating CIS^Sc^ recruitment to the membrane and subsequent firing.

Our work shows that CIS function in multicellular bacteria is distinct from Type 6 Secretion Systems and extracellular CIS, and possibly evolved due to the role CIS^Sc^ play in regulated cell death.

## Introduction

Bacteria employ different types of contractile injection systems (CIS) to mediate cell-cell interactions or cell death (1). CIS are evolutionarily and structurally related to contractile tails of bacteriophages and are comprised of core modules, including a baseplate, a contractile sheath, and an inner tube (2,3). CIS firing is triggered by a conformational change within the baseplate (4,5) and results in the contraction of the sheath, which in turn propels the inner tube along with the spike tip. Tube expulsion facilitates the release of associated effector proteins into target cells or the extracellular space (6,7).

Bioinformatic analyses have shown that CISs are conserved across diverse microbial phyla, including Gram-negative and Gram-positive bacteria, as well as archaea (8,9). Based on their distinct modes of action, CIS can be categorized into two main groups: intracellular type VI secretion systems (T6SS) and extracellular CIS (eCIS). T6SS are abundant among Gram-negative bacteria and in some archaea, and they are anchored to the cytoplasmic membrane during assembly and firing (10–12). A crucial component of the T6SS is the baseplate, which acts as a nucleus for T6SS assembly and mediates the binding of the T6SS particles to the cytoplasmic membrane until contraction. Upon contraction, the spike and inner tube are propelled out of the cell into an adjacent cell or the medium. eCIS, on the other hand, are assembled as free-floating particles in the cytoplasm and are subsequently released into the extracellular space upon lysis of the producer cell (13–16). Following the release, eCIS attach to the surface of target cells via their tail fibers, followed by contraction and puncturing of the target cell envelope.

Besides T6SS and eCIS, there is accumulating evidence of additional CIS assemblies with distinct modes of action in bacteria. For example, in multicellular cyanobacteria, CIS are anchored to the intracellular thylakoid membrane stacks via an extension of the baseplate components and have been proposed to function in cell lysis and the formation of “ghost cells” in response to stress conditions (17). Furthermore, several recent studies reported the production of free-floating cytoplasmic CIS particles in the multicellular monoderm *Streptomyces* bacteria, which have been shown to modulate cellular development through presumably slightly different mechanisms (18–22).

Notably, many *Streptomyces* genomes contain a class IId eCIS locus, a subtype that is structurally less well understood compared to other eCIS subtypes, such as ‘antifeeding prophages’ (AFPs) from *Serratia*, ‘metamorphosis-associated contractile structures’ (MACs) from *Pseudoalteromonas luteoviolacea*, AlgoCIS from *A. machipongonensis* or ‘*Photorhabdus* virulence cassettes’ (PVCs) from *P. asymbiotica* (1).

*Streptomyces* are best known for producing a plethora of medically and industrially important secondary metabolites, including molecules with antimicrobial, antifungal, anticancer or immunosuppressive properties. The production of these molecules is tightly coordinated with the *Streptomyces* developmental life cycle that encompasses functionally distinct filamentous cell types: vegetative hyphae that grow by tip extension and branching to scavenge for nutrients and reproductive (aerial) hyphae that eventually differentiate into chains of spores, which can then be dispersed to restart the life cycle (23,24). Using *Streptomyces coelicolor* as a model organism, we and others previously demonstrated that *S. coelicolor* CIS (CIS^Sc^) mediate a form of regulated cell death, which influences the onset of sporulation and secondary metabolite production in response to exogenous stress and unknown cellular signals (Figure 1a) (18,19).

**Figure 1:**
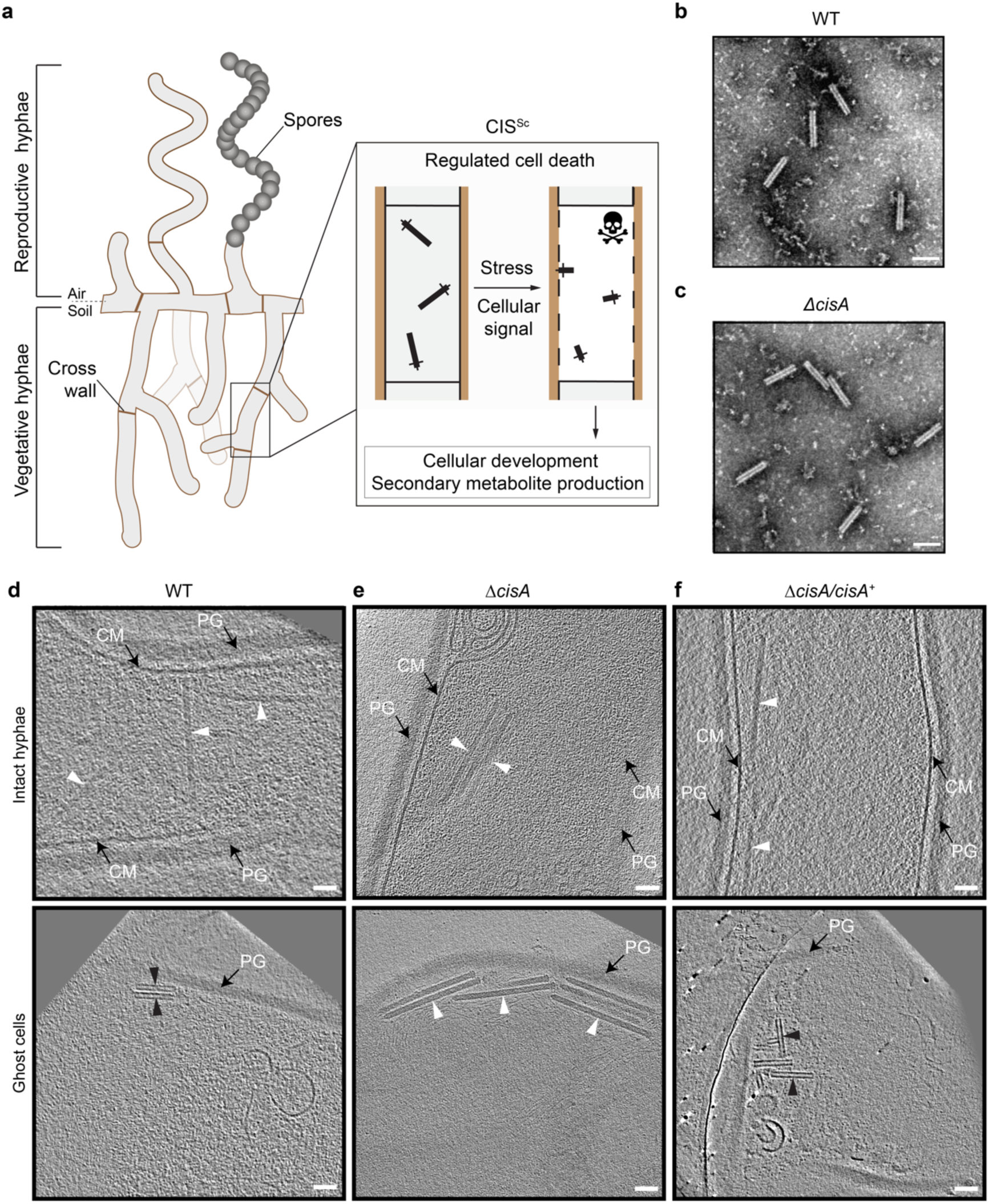
Contraction of CIS^Sc^ *in situ* is dependent on CisA. **a.** Schematic of the mode of action of CIS^Sc^ in *Streptomyces coelicolor* (18,19). CIS^Sc^ are assembled as free-floating particles in the cytoplasm of vegetative hyphae. In response to extracellular stress and/or an unknown cellular signal, CIS^Sc^ particles contract, which results in regulated cell death (mediated by released effectors) and impacts cellular development. **b/c.** Negative-stain electron micrographs of purified CIS^Sc^ particles from *S. coelicolor* wildtype (WT) (b) and the *ΔcisA* mutant (c), show that all CIS^Sc^ particles are functional and contract upon purification. Experiments were performed three independent times. Bars, 100 nm. **d-f.** Shown are representative images of cryo-electron tomogram slices (thickness 11 nm) of vegetative hyphae (top: intact cells; bottom: ghost cells) of *S. coelicolor* WT (d)*, ΔcisA* mutant (e), and the complemented *ΔcisA/cisA^+^* mutant (f). CIS^Sc^ particles remain almost exclusively in an extended state (white arrowheads) in the *ΔcisA* mutant, whereas in ghost cells derived from the WT and the complemented mutant, CIS^Sc^ particles (black arrowheads) are mostly contracted. See also Supplementary Fig. 1. PG, peptidoglycan; CM, cytoplasmic membrane; Bars, 50 nm.

It remains unclear, however, whether CIS^Sc^ contraction and firing occurs while the CIS^Sc^ particles are free-floating in the cytoplasm or whether this depends on an interaction with the membrane. Moreover, CIS encoded in *Streptomyces* genomes lack clear homologs for tail fiber components or a T6SS trans-envelope complex, raising the question of how CIS^Sc^ particles could potentially interact with the cytoplasmic membrane. We and others recently noted an uncharacterized protein (18,19), SCO4242, which is conserved in the majority of *Streptomyces* species and other Actinomycete species that carry a type IId eCIS locus (8,9) (Supplementary Fig. 1 and Supplementary Data 1 and 2). Interestingly, SCO4242 is encoded just downstream of the baseplate components in the CIS^Sc^ gene cluster, a location that usually encodes tail fibers from conventional eCIS (13,16,25). We will, therefore, refer to the gene product of SCO4242 as CisA for “CIS^Sc^-associated protein A.” Here, we set out to assess the importance of CisA for CIS^Sc^ function and its possible role as a membrane anchor.

## Results

### CisA is required for CIS^Sc^ contraction *in situ*

To explore the role of CisA (SCO4242) in CIS^Sc^ contraction and function, we first generated a *S. coelicolor ΔcisA* null mutant. Negative stain electron microscopy (EM) imaging of CIS^Sc^ particles that were purified from *ΔcisA* mutant cells showed assemblies in the contracted conformation, similar to the CIS^Sc^ particles purified from the wild type (WT) (Fig. 1b/c). Next, to examine the impact of CisA on the behavior of CIS^Sc^ *in situ*, we imaged vegetative hyphae of the WT, the *ΔcisA* mutant and the complemented mutant (Δ*cisA/cisA*^+^) by cryo-electron tomography (cryoET) (270 tomograms in total, n=3 experiments per strain). We previously demonstrated that CIS^Sc^ contraction correlates with the cellular integrity of *S. coelicolor* wild-type (WT) hyphae and showed that in intact hyphae, CIS^Sc^ particles are consistently observed in their extended conformation in the cytoplasm, whereas in partially lysed hyphae, the ratio of extended to contracted particles is approximately 2:1, and dead hyphae only contained fully contracted CIS^Sc^ assemblies (18). Interestingly, and in contrast to the WT and the complemented *cisA* mutant strain (Fig. 1d and f), *cisA-*deficient hyphae only contained CIS^Sc^ particles in the extended conformation irrespective of the cellular integrity of the hyphae (Fig. 1e, Supplementary Fig. 2a-d). Moreover, our analysis consistently showed only two CIS^Sc^ conformations in intact and dead hyphae, respectively: the fully extended state (233 nm in length, 18 nm in diameter, n=100 CIS^Sc^ particles) and the fully contracted state (124 nm in length, 23 nm in diameter, n=100 CIS^Sc^ particles). While hyphal cell death can be caused by a range of external or internal factors, our results demonstrate that CIS^Sc^ contraction is not a consequence of cell death and requires CisA *in situ*.

### CryoEM structure of the extended CIS^Sc^ assembly

The *S. coelicolor cis* gene cluster consists of 19 predicted open reading frames (accessions *SCO4242-4260*) (Fig. 2a), including genes with sequence similarities to potential CIS structural components (*cis1*-*cis16*), as well as additional genes with unknown functions (including *cisA*) (18,19). To determine whether CisA is an integral component of CIS^Sc^, we set out to elucidate the high-resolution structure of purified CIS^Sc^ particles from a non-contractile CIS^Sc^ mutant (18). The purified assemblies were subjected to single-particle cryo-electron microscopy (cryoEM) for structure determination (Supplementary Fig. 3), yielding the first high-resolution model of the CIS^Sc^ in its extended conformation, which also presents the first example of a CIS IId subtype (8).

**Figure 2:**
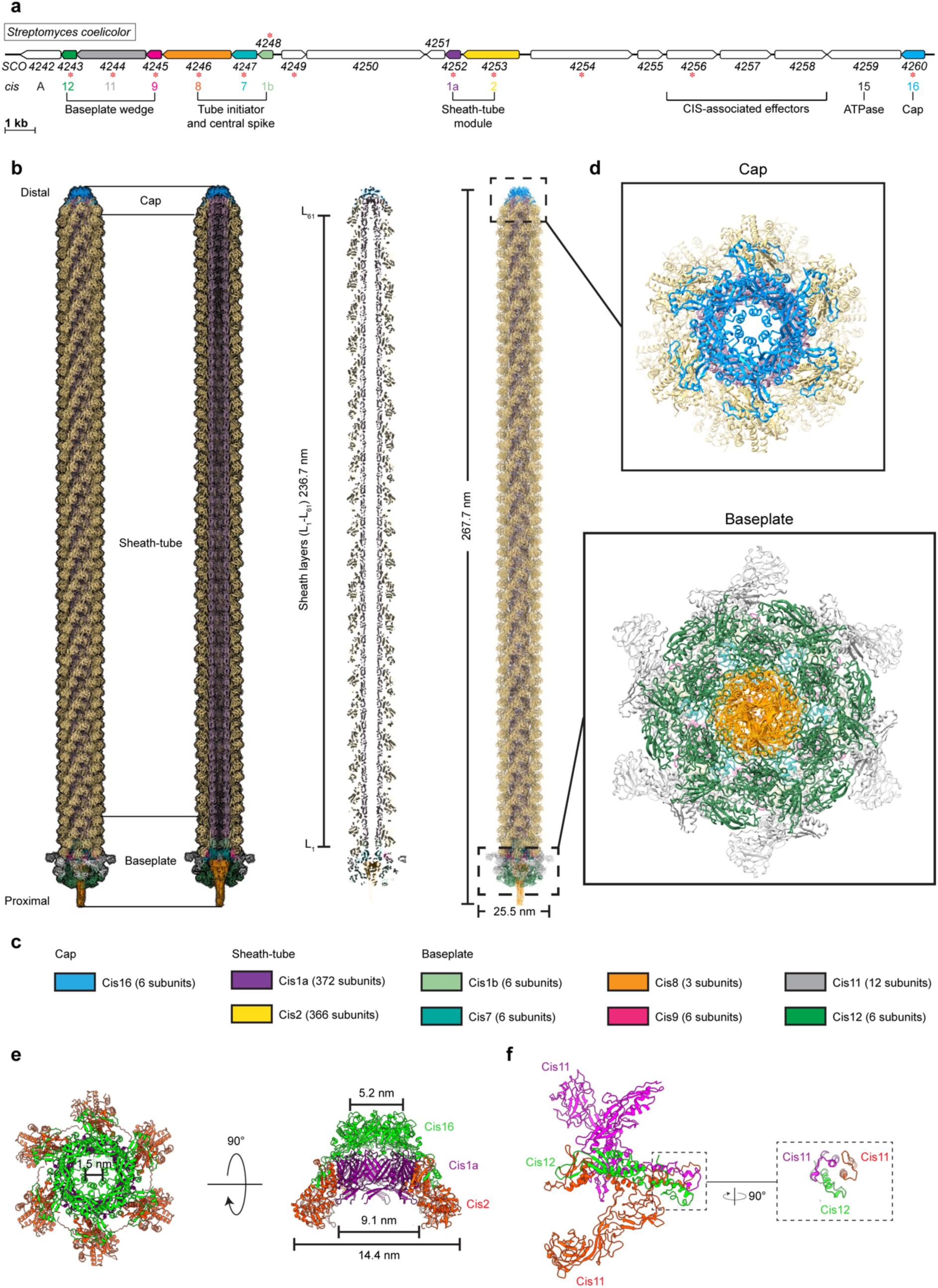
CryoEM structure of the extended CIS^Sc^ assembly reveals its composition. **a.** Schematic illustrating the CIS^Sc^ gene cluster (adapted from (18)). Asterisks indicate gene products that were detected by mass spectrometry analyzing crude preparations of CIS^Sc^ particles (red). See also Supplementary Table 1. **b/c.** Single-particle cryoEM structure of an extended CIS^Sc^ assembly obtained purified from a non-contractile CIS^Sc^ mutant. Shown are the composite atomic model in surface (left) and ribbon (right) rendering (b). Subunits are color-coded according to (a) and their copy numbers in the assembly are indicated in (c). **d.** Perpendicular views of the ribbon representation of the CIS^Sc^ model of the cap and baseplate modules. **e.** Top view (left) and side view (right) of ribbon diagrams showing the cap module of CIS^Sc^, revealing that the cap is composed of a Cis16 hexamer. **f.** Ribbon diagrams showing a single baseplate wedge subunit, which is composed of two copies of Cis11 and one copy of Cis12.

Our model shows that extended CIS^Sc^ particles share the conserved eCIS architecture, including the three core modules: cap, sheath-tube and baseplate (Fig. 2b). We processed the dataset using an established workflow that was previously employed to produce the atomic model for other CIS (Supplementary Fig. 3) (13,17). The resulting maps of the baseplate and the cap reached resolutions of 3.5 and 3.4 Å, respectively. The quality of the final maps from the two modules enabled *de novo* structural modeling (Supplementary Fig. 4, Supplementary Table 1).

The final model of CIS^Sc^ comprises 783 polypeptide chains that assemble into a bullet-shaped particle measuring 268 nm in length (Fig. 2b/c). All analyzed particles were identical in length, consistent with our *in situ* data of extended CIS^Sc^ particles in vegetative hyphae of WT *S. coelicolor* (Fig. 1d-f). The length of eCIS particles is often controlled by tape measure proteins (26). We note, however, that the corresponding gene is absent from the CIS^Sc^ gene cluster. The distal end of the CIS^Sc^ is capped by a tail terminator complex (Fig. 2c/e). This cap comprises one protein, Cis16, forming a C6-symmetrical complex that terminates the inner tube and sheath, respectively. This minimal architecture resembles the cap complexes of PVC or Pyocin R2 CISs, but it is different from the more complex AlgoCIS from *A. machipongonensis* and AFP from *S. entomophila* (Supplementary Fig. 5) (13,16,25,27). The structures of the CIS^Sc^ sheath-tube module have been previously discussed, notably showing that the sheath (domains 1/2) contracts by widening and shortening, similar to other CIS (18). We found that the module consists of 61 sheath layers (Cis2-L_1_–L_61_) and 60 tube layers (Cis1-L_1_-L_60_) (Fig. 2b/d). The baseplate module comprises a heteromeric assembly of the proteins Cis1b/7/8/9/11/12 (Fig. 2b/c). The symmetry transition from the inner tube (Cis1a, C6) to the VgrG-like spike (Cis8, C3) is adapted by two layers of the tube initiators Cis1b and Cis7. The first layer of the sheath (Cis2) is bound to the conserved gp25-like protein Cis9, which connects to the baseplate ‘wedges’ composed of Cis11 and Cis12 (Fig. 2b/c). Importantly, the CIS^Sc^ ‘wedges’ consist of two copies of Cis11 and one copy of Cis12 (Fig. 2f), resembling the baseplate of Pyocin R2(27). This is in contrast to other previously reported eCIS, whose baseplate wedge comprises only one Cis11/12 component (AlgoCIS, AFP, PVC) and a Cis11 extension forming a cage around the spike (AlgoCIS) (Supplementary Fig. 6) (13,16,25). Furthermore, we did not detect any structural components that resemble tail fibre proteins. This is in line with the bioinformatic analysis of the *cis* gene cluster (Fig. 2a), suggesting the absence of conventional tail fibre genes.

Out of the 19 predicted open reading frames in the CIS^Sc^ gene cluster (Fig. 2a), nine encoded proteins were present in the final reconstruction, for which all atomic models were built (Fig. 2b/c). However, we did not detect a density for CisA in our model, suggesting that CisA is not part of purified CIS^Sc^ particles but may interact with them via a different mechanism. This finding is also in agreement with mass spectrometry results, which indicate the absence of CisA peptides from purified CIS^Sc^ particles (Supplementary Table 1) (18). We speculate that the purification protocol used to isolate CIS^Sc^ particles for single-particle analysis may not preserve the CIS^Sc^-CisA interaction or that the interaction is too transient and only occurs under specific growth conditions. To better understand how CisA mediates CIS^Sc^ function, we further characterized CisA *in vivo*.

### CisA is a bitopic protein

Bioinformatic analyses (18,19) and structural modeling of CisA suggest that CisA consists of a largely unstructured N-terminal region, a transmembrane segment, and a conserved C-terminal domain (CTD) that shares structural similarity to periplasmic substrate-binding domains (Fig. 3a). Additional Alphafold3 (28) modeling using just the ordered C-terminus of CisA, including the transmembrane domain and the CTD (amino acids 285-468), predicts that CisA can oligomerize into a pentamer via the CTD (Fig. 3b and c). These findings suggest that CisA may exist in distinct configurations within the cell. To characterize the intracellular localization of CisA, we first generated a strain in which *cisA* was fused to the N-terminus of *mCherry* and expressed *in trans* from a constitutive promoter in the *S. coelicolor* Δ*cisA* mutant. Using fluorescence light microscopy, we consistently observed a CisA-mCherry fluorescence signal along the hyphal periphery, indicative of a membrane-associated protein (Fig. 3d). In contrast, no fluorescence was observed in WT hyphae carrying an empty plasmid (Fig. 3e). Next, we tested if CisA is indeed localized to the membrane *in vivo* by performing cellular fractionation experiments. We separated soluble proteins from membrane proteins using whole cell lysates from the *S. coelicolor* WT or the Δ*cisA* mutant that was complemented with a *cisA-3xFLAG* fusion expressed *in trans* from a constitutive promoter. Our analysis confirmed that CisA-3xFLAG co-sedimented with the cell membranes, while the cytoplasmic transcription factor WhiA could only be detected in the soluble protein fraction (Fig. 3f).

**Figure 3:**
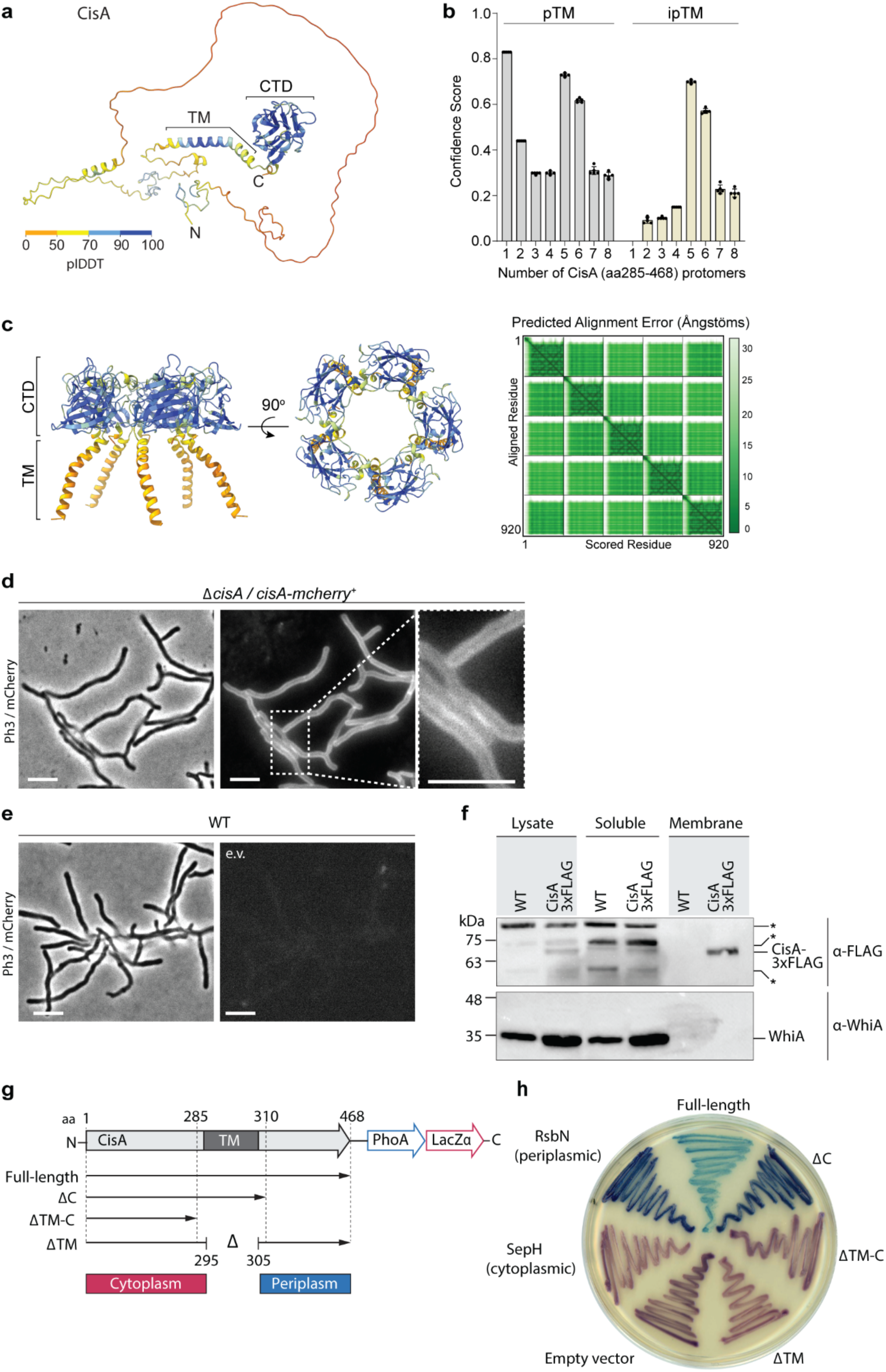
CisA is a single-pass membrane protein predicted to oligomerize. **a.** Alphafold3 model of CisA colored in pLDDT values. CisA is predicted to contain a largely unstructured N-terminal domain, a transmembrane domain (TM) and a conserved C-terminal (CTD). **b.** Confidence scores for the predicted oligomeric configuration of CisA. Shown are the predicted template modeling (pTM) and the predicted interface pTM scores (ipTM) obtained from *n* = 5 predictions per CisA configuration. An ipTM score greater than 0.7 indicates a likely protein–protein interaction. CisA aa 285-468, including the transmembrane domain and the CTD (amino acid 285-468), was used as input sequences for modeling. **c.** Alphafold3 model of a CisA (amino acids 285-468) pentamer and the corresponding predicted alignment error (PAE) plot. **d/e.** Representative micrographs (left panels: phase contrast, Ph3; right panel: mCherry) of strains of *S. coelicolor* vegetative hyphae either constitutively expressing CisA*-*mCherry (Δ*cisA/cisA-mcherry*) *in trans* (d) or WT cells carrying an empty vector (e.v.) to control for background fluorescence (e). Box shows a magnified region of hyphae with CisA*-*mCherry accumulation in the cellular periphery. Bars, 5 µm. **f.** Western blot of the cellular fractionation of samples from the *S. coelicolor* WT or the *ΔcisA* mutant constitutively expressing a CisA-3xFLAG fusion *in trans*. Lysate and soluble and membrane fractions were probed for the presence of CisA-3xFLAG with an α-FLAG antibody (top). Fractionation efficiency was assessed using an α-WhiA antibody to detect the soluble transcription factor WhiA (bottom). The same volumes for each fraction were loaded and analyzed by immunoblotting. CisA-3xFLAG was detected in the lysate and the membrane fraction. Asterisks denote non-specific signals. The experiment was performed in biological duplicates and shown is a representative image. **g/h.** Experimental determination of the CisA membrane topology using the dual *phoA*-*lacZα* reporter in *E. coli*. The schematic diagram (g) depicts the four CisA constructs used in the assay and indicates the relevant amino acid (aa) deleted in the three CisA mutant derivates. The analysis of colony coloration (h); pink coloration for cytosolic proteins and blue coloration for periplasmic proteins, indicates that the CisA N-terminus is localized in the cytoplasm and the CisA C-terminus localized to the periplasm and this topology is dependent on the transmembrane domain (TM). *E. coli* strains carrying an empty reporter plasmid or reporter plasmids with the *Streptomyces* genes for SepH (cytoplasmic) and RsbN (periplasmic) were used as controls. Note that the expression of full-length CisA in *E. coli* is toxic, resulting in reduced growth and a lighter blue coloration.

Next, we investigated the topology of CisA in the membrane using a dual *phoA*-*lacZ*α reporter system in *E. coli*. The reporter activity can be directly visualized on indicator plates based on the complementary activities of the cytoplasmic reporter β-galactosidase LacZ (magenta coloration) and the periplasmic reporter alkaline phosphatase PhoA (blue coloration) (29). We engineered *E. coli* strains that expressed a C-terminal fusion of CisA variants to the PhoA-LacZα reporter, including full-length CisA and CisA variants that carried either a deletion in the predicted transmembrane domain and/or the putative periplasmic domain (Fig. 3g). In addition, we included three controls, including 1) cells carrying the empty reporter plasmid, resulting only in cytoplasmic β-galactosidase activity, 2) cells expressing a PhoA/LacZ fusion to the *Streptomyces* cytoplasmic cell division protein SepH, and 3) cells expressing a PhoA/LacZ fusion to the periplasmic domain of the *Streptomyces* membrane protein RsbN (30,31). The results of this assay confirmed that CisA is a bitopic protein with an N-terminus (amino acids 1-285) located in the cytoplasm and a C-terminus (amino acids 310-468) present in the periplasm of *E. coli* (Fig. 3h).

Collectively, our results demonstrate that CisA is a membrane protein comprised of a flexible cytoplasmic N-terminal domain and a periplasmic C-terminal domain that is predicted to facilitate CisA oligomerization.

### CisA is essential for the cellular function of CIS^Sc^

We previously established a fluorescence-based assay to show that the production of functional CIS^Sc^ particles results in a significantly reduced viability of *Streptomyces* hyphae exposed to exogenous stress, including treatment with the membrane-pore forming antibiotic nisin, UV radiation or the protonophore carbonyl cyanide 3-chlorophenylhydrazone (CCCP) (18). This viability assay is based on the measurement of the ratio of two fluorescent markers, namely cytoplasmic sfGFP produced by viable intact hyphae and the membrane dye FM5-95 to detect intact as well as partially lysed hyphae. Using this assay, we tested whether CisA was essential for CIS^Sc^-mediated cell death and determined the relative viability of the WT, the Δ*cisA* and the complemented *cisA* (Δ*cisA/cisA*^+^) strains expressing cytosolic sfGFP. We compared cells exposed to a sublethal concentration of nisin (1 µg/ml for 90 min) to cells not exposed to antibiotic stress. All three strains were found to express CIS^Sc^ assemblies under these growth conditions (Supplementary Fig. 7).

Without nisin stress, all three strains exhibited a similar sfGFP/FM5-95 ratio, indicating no significant difference in viability (Fig. 4a/c). As previously observed (18), the viability of the WT was dramatically reduced in response to nisin stress. In contrast, the viability of Δ*cisA* hyphae was unaffected by nisin stress and comparable to the viability of untreated cells. This phenotype could be completely reversed by complementing the Δ*cisA* deletion *in trans* (Fig. 4b/d). Notably, the Δ*cisA* strain phenocopies CIS^Sc^ knockout and non-contractile mutants we analyzed previously (18). Thus, we conclude that CisA is required for CIS^Sc^-mediated cell death upon stress.

**Figure 4:**
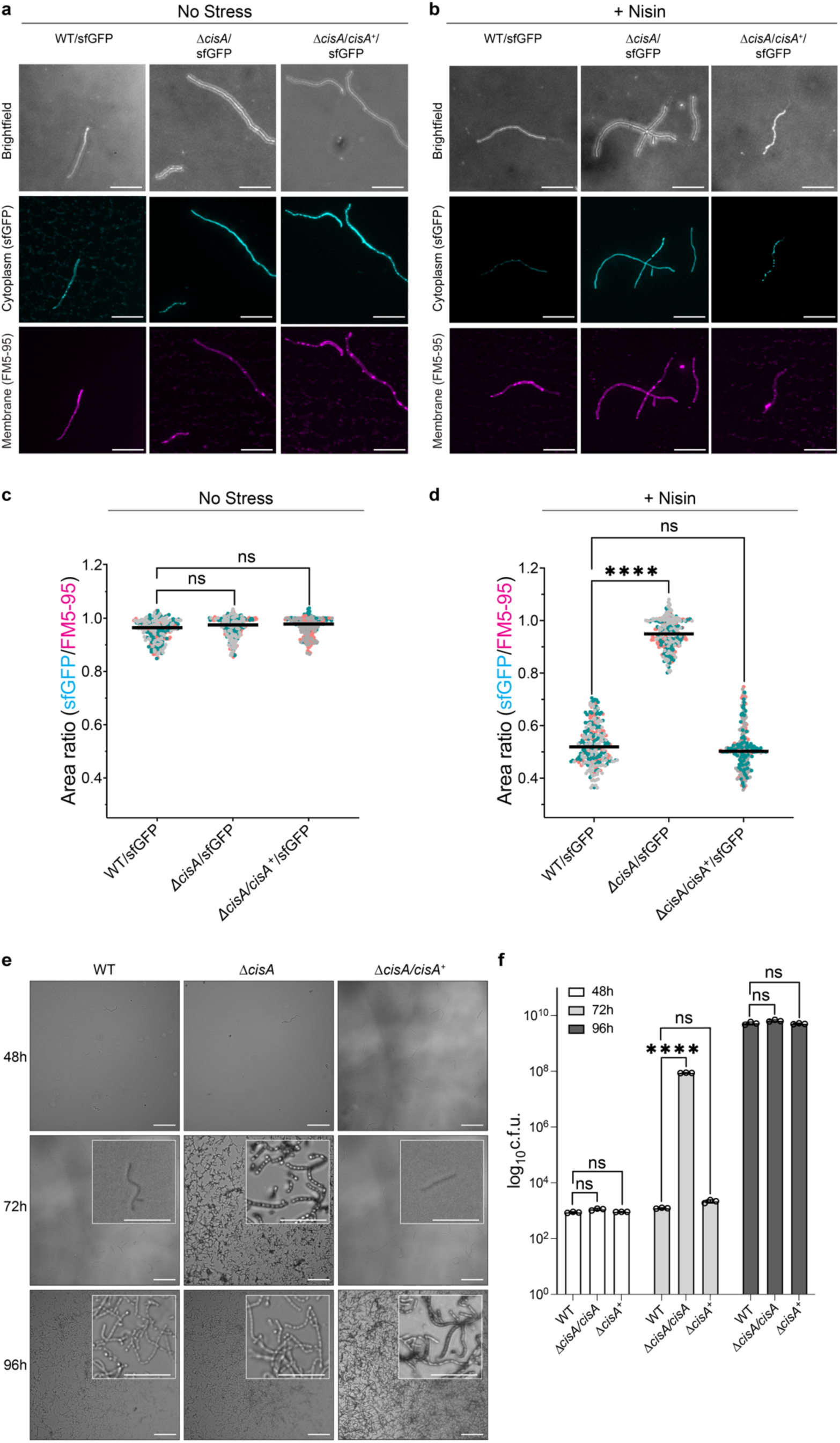
CisA is required for regulated cell death and cellular function of CIS^Sc^. **a-d.** Light microscopy-based viability assay showing that CIS^Sc^ do not mediate cell death in *cisA*-deficient hyphae in response to exogenous stress. Shown are representative micrographs of the WT/sfGFP, the *ΔcisA*/sfGFP and the complemented *ΔcisA/cisA^+^*/sfGFP mutant expressing cytosolic sfGFP (indicator for live cells). Strains were grown for 48 h and then either incubated without (a) or with (b) the membrane-disrupting antibiotic nisin (1 µg/ml nisin for 90 min). Samples were briefly vortexed to break up mycelial clumps and subsequently stained with the fluorescent membrane dye FM5-95 to stain cells with an intact or damaged cytoplasmic membrane. Bars, 10 µm. Z-stacks were taken of the samples, and the resulting relative fluorescence ratios of sfGFP to FM5-95 are shown in c/d. WT and *ΔcisA/cisA^+^* complemented cells showed a significantly higher rate of cell death in response to nisin treatment than *ΔcisA* cells. There is no significant difference in the viability of the tested strains in the absence of nisin. Experiments were performed in three biological replicates, as indicated by the green, grey, and magenta data points. Black lines indicate the mean ratio derived from biological triplicate experiments (n=100 images per replicate). ns (not significant) and **** (p < 0.0001) were determined using a one-way ANOVA and Tukey’s post-test. **e.** Visualization of spore production in WT, *ΔcisA* mutant and *ΔcisA/cisA^+^*strains reveal accelerated cellular development and sporulation of the *ΔcisA* mutant. Shown are representative brightfield images of surface impressions of plate-grown colonies of each strain taken at the indicated time points. Only sporulating hyphae will attach to the hydrophobic cover glass surface. Insets show magnified regions of the colony surface containing spores and spore chains. Bars, 50 µm. **f.** Quantification of sporulation shown in (e) via counts of colony-forming units (c.f.u.). The graph shows a 6-fold higher c.f.u. count at 72 h in the *ΔcisA* mutant. Strains were grown on R2YE agar and spores were harvested after 48 h, 72 h, and 96 h of incubation. Data show mean ± s.d. obtained from biological triplicate experiments. ns (not significant) and **** (p < 0.0001) were determined using a one-way ANOVA and Tukey’s post-test.

To further test the dependence of CIS^Sc^ function on CisA, we conducted additional *in vivo* assays. Previous studies showed that the absence of functional CIS^Sc^ impacted the timely differentiation of *Streptomyces* hyphae into chains of spores, resulting in accelerated cellular development and reduced secondary metabolite production (18,19). Here, we repeated these analyses using the WT, the Δ*cisA* mutant and the complemented mutant (Δ*cisA/cisA^+^*). All strains consistently completed their life cycle and synthesized spores after 96 h of growth on solid medium (Fig. 4e). Importantly, unlike the WT and the Δ*cisA/cisA^+^* strain, Δ*cisA* mutant colonies sporulated markedly earlier (72 h). These findings were further corroborated by quantifying the number of spores produced by each strain under the same experimental conditions (Fig. 4f). Finally, we examined the effect of a *cisA* deletion on secondary metabolite production by quantifying the level of actinorhodin production in *S. coelicolor* (32). Compared to the WT and the complemented mutant strain (*ΔcisA/cisA^+^*), the Δ*cisA* mutant showed significantly reduced levels of actinorhodin production (Supplementary Fig. 8). Our data therefore support the conclusion that the cellular function of CIS^Sc^ in *S. coelicolor* depends on CisA, and its absence impacts the timely cellular and chemical differentiation of *S. coelicolor* hyphae.

## Discussion

Here, we identify CisA as an essential membrane-associated factor that mediates the cellular function of CIS^Sc^ particles in *Streptomyces coelicolor*. In the absence of CisA, CIS^Sc^ are not able to contract in the cellular context (Fig. 1). Similar to a CIS^Sc^ deletion or non-contractile mutant (18), the deletion of *cisA* decreases the induction of cell death upon stress, which in turn affects cellular development and secondary metabolite production (Fig. 5). The mechanistic aspects of this distinct mode of action of CIS^Sc^ are discussed below.

**Figure 5:**
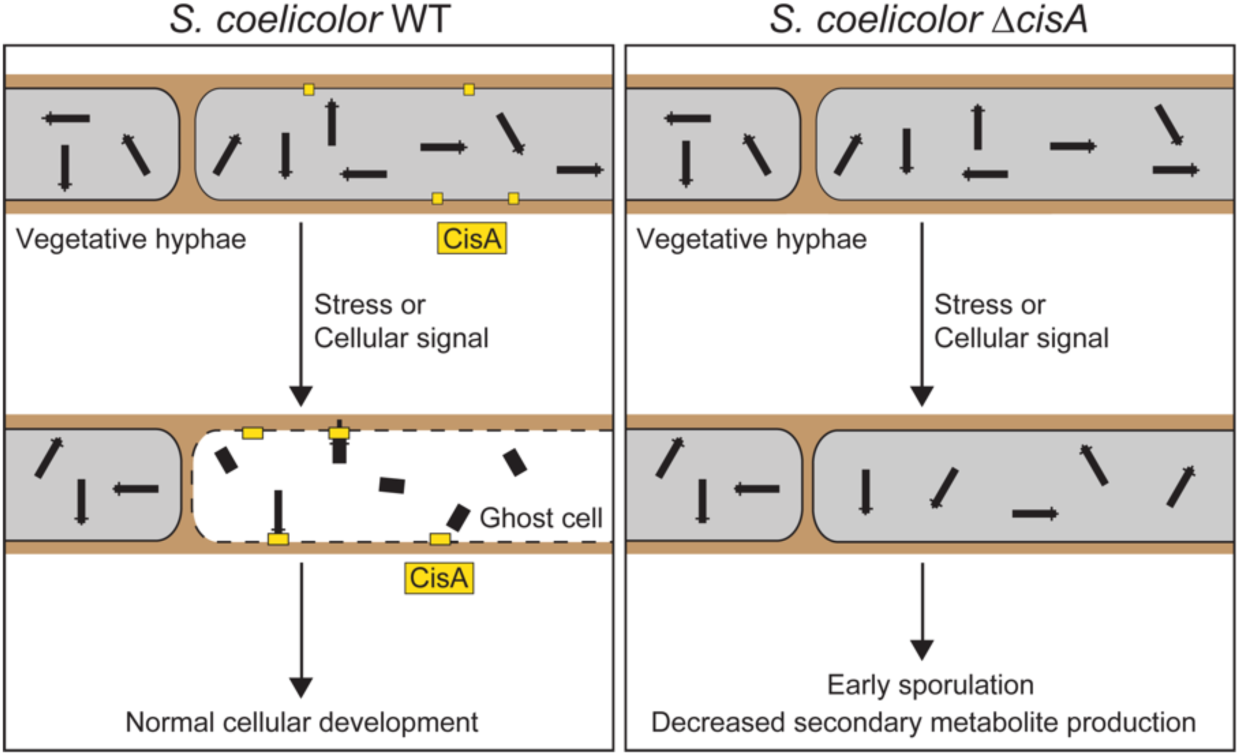
Cytoplasmic CIS^Sc^ require the membrane protein CisA for contraction and function. Proposed model illustrating a possible role of CisA. In response to exogenous stress or an unknown cellular signal, membrane-bound CisA either directly or indirectly mediates the association of free-floating CIS^Sc^ particles with the cytoplasmic membrane, leading to CIS^Sc^ firing and regulated cell death, which impacts the *Streptomyces* developmental life cycle.

A conserved feature of all previously studied CIS is the requirement of the contractile apparatus to bind to a membrane before firing. This can be achieved in various ways, resulting in the classification of CIS into distinct modes of action. First, T6SS are anchored to the cytoplasmic membrane of the host cell by a trans-envelope complex spanning from the cytoplasm to the inner membrane, periplasm, and outer membrane (33–35). Second, contractile phages and eCIS bind to the surface of their target cell via tail fibers, which are typically connected to the baseplate and transmit the signal for firing to the baseplate (13,16,36). Third, tCIS are anchored to thylakoid membrane stacks in cyanobacteria by an extension of their baseplate (17). Finally, and in contrast to the previously described CIS, we demonstrate that intracellular CIS^Sc^ particles from *Streptomyces* do not contain structural components that can directly bind to the host membrane. Our data suggest that CisA is required to mediate an interaction of CIS^Sc^ with the cytoplasmic membrane prior to firing (Fig. 5). This interaction may occur directly via the binding of the CIS^Sc^ particle to the membrane protein CisA, or through the interaction with a yet unknown factor. To explore this idea, we performed an *in silico* prediction of protein-protein interactions between monomeric CisA and CIS^Sc^ components using Alphafold2-Multimer (37) (Supplementary Fig. 9a). Interestingly, this analysis identified the baseplate component Cis11 as a significant hit and possible interaction partner of CisA (Supplementary Fig. 9b-d). Importantly, such a protein-protein interaction would be consistent with our cryoEM structure (Fig. 2b-f), which shows a peripheral surface-exposed position of Cis11 in the baseplate complex of extended CIS^Sc^.

Additional structural modeling with truncated versions of CisA in complex with Cis11 further support the idea that the largely unstructured cytosolic portion of CisA is required to interact with the Cis11 (Supplementary Fig. 10). We have been unable to confirm a direct interaction between CisA and Cis11 because the expression of *cisA* is toxic in *E. coli,* which prevented the experimental analysis of the CisA-Cis11 interaction using co-purification approaches and bacterial two-hybrid studies. While we currently cannot exclude the possibility that the proposed CisA-Cis11 interaction involves an additional unknown factor, we did detect CisA peptides in crude purifications of CIS^Sc^ from nisin-stressed cells (Supplementary Table 2). This would be consistent with a transient and/or short-lived interaction of CIS^Sc^ particles with the membrane and CisA. In agreement with this idea, we rarely observed extended CIS^Sc^ assemblies associated with the cytoplasmic membrane via their baseplate in WT cells (Fig. 1d).

We further speculate that CisA may not only serve as a mediator for the interaction of CIS^Sc^ with the cytoplasmic membrane but also as a sensor and checkpoint for inducing cell death under specific stress conditions or in response to cellular signals. Notably, protein structure searches with Foldseek (38) using just the conserved extra-cytoplasmic C-terminal CisA domain (amino acids 310-468) suggest that this domain shares similarity with protein solute-binding protein domains. Thus, the C-terminal CisA domain could potentially be involved in directly or indirectly sensing stress signals. Recognition or binding of such a signal by monomeric CisA could trigger a conformational change that results in CisA oligomerization, which in turn could lead to an “activated” CisA configuration that is primed to mediate the recruitment of CIS^Sc^ assemblies to the cytoplasmic membrane, followed by firing.

It is conceivable that the lethal consequences of CIS^Sc^ firing for the producing cell may have driven the evolution of a stepwise mechanism, involving a checkpoint that prevents self-inflicted cell death by accidental firing. Such a stepwise mechanism would ensure that the CIS^Sc^ remain in a free-floating, and safe state within the cytoplasm until a stress signal induces CisA-mediated CIS^Sc^ contraction.

## METHODS

### Bacterial strains, plasmids, and oligonucleotides

Bacterial strains, plasmids, and oligonucleotides used in this study are listed in Supplementary Tables 3-4. Plasmids were generated using either standard restriction-ligation or assembled using the Gibson Assembly Master Mix (NEB) or *In Vivo* Assembly. All plasmids were verified by DNA sequencing. *Escherichia coli* strains were cultured in LB, SOB, or DNA medium. *E. coli* strains TOP10, DH5α and NEB5α were used to propagate plasmids and cosmids, *E. coli* strain BW25113/pIJ790 for recombineering cosmids and *E. coli* ET12567/pUZ8002 for interspecies conjugation. When required, media was supplemented with antibiotics at the following concentrations: 100 µg/ml carbenicillin, 50 µg/ml apramycin, 50 µg/ml kanamycin, 50 µg/ml hygromycin.

*Streptomyces coelicolor* strains were cultivated in LB, TSB, TSB-YEME, or R2YE liquid medium at 30 °C in baffled flasks or flasks with springs, at 250 rpm or grown on LB, SFM, R2YE medium solidified with 1.5 % (w/v) Difco agar (39). Antibiotics were added at the following concentrations: 25 µg/ml apramycin, 50 µg/ml kanamycin, 25 µg/ml hygromycin, 12.5 µg/ml nalidixic acid.

### Construction of the *S. coelicolor* Δ*cisA* mutant

The λ RED homologous recombination system was used to isolate gene replacement mutations using PCR-directed mutagenesis (ReDirect) of the *S. coelicolor* cosmid StD8A containing the *cis* gene cluster(40),(41). The *cisA* coding sequence (*SCO4242*) was replaced with the *aac3(IV)-oriT* resistance cassette from pIJ773. The mutant cosmid (pSS684) was introduced into *E. coli* ET12567/pUZ8002, followed by conjugation into *S. coelicolor* M145. Exconjugants that had successfully undergone double-homologous recombination were identified by screening for apramycin resistance and kanamycin sensitivity. The deletion of *cisA* was subsequently verified by PCR.

### Sheath preparation of CIS^Sc^

*S. coelicolor* strains were grown in 30 ml TSB, TSB-YEME or R2YE liquid medium for 48 h. Cells were pelleted by centrifugation (7,000 x g, 10 min, 4 °C), resuspended in 5 ml lysis buffer (150 mM NaCl, 50 mM Tris-HCl, 0.5× CellLytic B (Sigma-Aldrich), 1 % Triton X 100, 200 µg/ml lysozyme, 50 μg/ml DNAse I, pH 7.4), and incubated for 1 h at 37 °C. Cell debris was removed by centrifugation (15,000 × g, 15 min, 4 °C) and cleared lysates were subjected to ultra-centrifugation (150,000 × g, 1 h, 4 °C). Pellets were resuspended in 150 µl resuspension buffer (150 mM NaCl, 50 mM Tris-HCl, supplemented with protease inhibitor cocktail [Roche], pH 7.4). CIS^Sc^ sheath preparations were analyzed by negative stain EM imaging (42) and mass spectrometry at the Functional Genomics Center Zürich.

To purify non-contractile CIS^Sc^ particles from *S. coelicolor (*SS393), cleared cell lysates were subjected to ultracentrifugation using a sucrose cushion (20 mM Tris pH 8.0, 150 mM NaCl, 50 mM EDTA, 1 % Triton X 100, 50 % [w/v] sucrose) (18). The sucrose cushion alongside 1 mm of liquid above and residual bacterial contaminants were removed by centrifugation at 15,000 x g. Samples were then subjected to a second round of ultracentrifugation without a sucrose cushion (150,000 ξ g, 1 h, 4 °C) and resulting cell pellets were resuspended in buffer (50 mM Tris, 150 mM NaCl). These crude samples were then further purified by gradient ultracentrifugation (10-50 % continuous gradient made with BIOCOMP gradient master IP model 107 gradient forming instrument) at 100,000 x g for 1 h using a SW55 Ti rotor. The gradient was analyzed in 11 fractions of 500 µl, which were imaged by negative stain EM. The fractions, which contained CIS^Sc^ particles, were pooled and used for further experiments.

### Negative stain electron microscopy

4 µl of purified CIS^Sc^ sheath particles were adsorbed to glow-discharged, carbon-coated copper grids (Electron Microscopy Sciences) for 60 s, washed twice with milli-Q water and stained with 2 % phosphotungstic acid for 45 s. The grids were imaged at room temperature using a Thermo Fisher Scientific Morgagni transmission electron microscope (TEM) operated at 80 kV.

### Mass spectrometry analysis

To confirm the presence of predicted CIS^Sc^ components, isolated sheath particles were subjected to liquid chromatography–mass spectrometry analysis (LC–MS/MS). These experiments were conducted at the Functional Genomics Center Zürich. First, the samples were digested with 5 µl of trypsin (100 ng/µl in 10 mM HCl) and microwaved for 30 min at 60 °C. The samples were then dried, dissolved in 20 µl ddH_2_0 with 0.1 % formic acid, diluted in 1:10 and transferred to autosampler vials for liquid chromatography with tandem mass spectrometry analysis. A total of 1 µl was injected on a nanoAcquity UPLC coupled to a Q-Exactive mass spectrometer (ThermoFisher). Database searches were performed by using the Mascot swissprot and tremble_streptomycetes search programs. For search results, stringent settings have been applied in Scaffold (1 % protein false discovery rate, a minimum of two peptides per protein, 0.1 % peptide false discovery rate). The results were visualized by Scaffold software ([Proteome Software Inc.], Version 4.11.1).

### Protein gel electrophoresis and western blotting

For general protein analysis, protein samples were boiled at 95 °C for 5 min and separated by a 4–20% Mini-PROTEAN® TGX Stain-Free™ SDS PAGE (BioRad) or 12 % Tris-Glycine SDS PAGE (Invitrogen) and visualized using InstantBlue Coomassie Protein Stain or ReadyBlue Protein Gel Stain (Sigma Aldrich).

For western blotting, cells were grown in biological triplicate for 48 h in TSB, 2 ml aliquots of each culture were pelleted and washed once with 1x PBS. Cell pellets were resuspended in 0.4 ml of sonication buffer (20 mM Tris pH 8.0, 5 mM EDTA, 1x EDTA-free protease inhibitors [Sigma Aldrich]) and subjected to sonication at 4.5-micron amplitude for 7 cycles of 15 s on / 15 s off. Samples were centrifuged at 17,000 x *g* for 15 min at 4 °C. Protein concentration in cleared cell lysate supernatants was determined by Bradford Assay (Biorad). Equivalent total protein concentrations (1 mg/ml) were assayed using the semi-dry western blot transfer. The gel was transferred to a PVDF membrane (Biorad) and probed with the appropriate antibody diluted in TBS-T or using the iBIND buffer system (Thermo Fisher): Rabbit anti-FLAG ([Sigma Aldrich] F7425, 1:2500), Rabbit anti-WhiA ((43), 1:2500), and Goat Anti-Rabbit-HRP (Abcam Ab6721, 1:5000). To visualize HRP-conjugated antibodies, membranes were incubated with SuperSignal West Femto ECL solution (Thermo Fisher).

### Cellular fractionation

To fractionate membrane and soluble proteins from *S. coelicolor* (WT, SS576), cells were grown for 48 h in 50 ml TSB/YEME and the entire culture volume was pelleted by centrifugation. Cell pellets were resuspended in 1/10 volume of lysis buffer (0.2 M Tris-HCl, pH 8, 10 mg/mL lysozyme, and 1× EDTA-free protease inhibitors; [Roche]), incubated for 30 min at 37 °C, and then briefly cooled on ice before lysed by sonication (11 x 15 sec on/off at 50 % power at 8 microns on ice). Cell debris was removed by centrifugation at 16,000 × g for 20 min and the supernatant was subsequently subjected to two additional rounds of centrifugation. The cleared cell lysate was subjected to ultracentrifugation for 1 h at 150,000 × g at 4 °C to separate the soluble proteins from membrane proteins. Sedimented membrane proteins were resuspended and washed twice in 1 volume of wash buffer (60 mM Tris-HCl, 200 mM NaCl, pH 8, 0.2 mM EDTA, and 0.2 M sucrose) at 150,000 × g at 4 °C for 1 h. The wash step was repeated one final time with the wash buffer containing 8 M urea to remove traces of membrane-associated proteins (44). The final pellet was dissolved in 1/10 of the initial volume with wash buffer (no urea). Equi-volume amounts of fractions were mixed with 2x SDS sample buffer and analyzed by immunoblotting.

### Membrane protein topology analysis in *E. coli*

The coding sequence of CisA was inserted in the dual pho-lac reporter plasmid pKTop, which consists of an *E. coli* alkaline phosphatase fragment PhoA_22-472_ fused in frame after the α-fragment of β-galactosidase LacZ_4-60_ (Supplementary Table 3) (29). For membrane protein topology, *E. coli* TG1 was transformed with the pKTop and derivatives. Membrane protein topology was assayed by plating the resulting reporter strains on dual-indicator plates containing LB agar supplemented with 80 μg/ml 5-bromo-4-chloro-3-indolyl phosphate disodium salt (X-Phos) (Sigma Aldrich, RES1364C-A101X) and 100 μg/mL 6-chloro-3-indolyl-β-d-galactoside (Red-Gal) (Sigma Aldrich, B6149) as indicators, 1 mM IPTG, and 50 μg/ml kanamycin. A periplasmic or extracellular location of the reporter fusion results in higher alkaline phosphatase activity (blue color), whereas a cytosolic location of the reporter leads to higher β-galactosidase activity (magenta color). Plates were incubated for two days at 37 °C and then scanned.

### Vitrified sample preparations

For single-particle cryoEM (SPA), the *S. coelicolor* non-contractile CIS particles were purified as described above and vitrified using a Vitrobot Mark IV (Thermo Fisher Scientific). 4 µl of samples were applied twice on glow-discharged 200 mesh Quantifoil copper grids (R 2/2) which were manually coated with a layer of 1 nm carbon. Grids were blotted for 5.5 s and plunge-frozen in liquid ethane-propane mix (37 %/63 %). Frozen grids were stored in liquid nitrogen until loading into the microscope.

For cryo-electron tomography (cryoET), *Streptomyces* cells were mixed with 10 nm Protein A conjugated colloidal gold particles (1:10 v/v, [Cytodiagnostics]) and 4 µl of the mixture was applied to a glow-discharged holey-carbon copper EM grid (R2/1 or R2/2, [Quantifoil]). The grid was automatically blotted from the backside for 4-6 s in a Mark IV Vitrobot by using a

Teflon sheet on the front pad, and plunge-frozen in a liquid ethane-propane mixture (37 %/63 %) cooled by a liquid nitrogen bath.

### SPA data collection and image processing

44,925 movies were collected on Titan Krios microscope (Thermo Fisher Scientific), operated at 300 keV equipped with K3 direct electron detector, operating in counting mode, and using a slit width of 20 eV on a GIF-Quantum energy filter (Gatan). The automated data collection was conducted with EPU software (Thermo Fisher Scientific) with a final pixel size of 1.065Å/pix over 40 frames with a total dose of 50 e^-^/Å^2^. The targeted defocus was set between −1.6 and −2.8 µm with 0.2 µm increment.

Movie-alignments with dose-weighting were performed in MotionCor2 software (45) with a gain reference estimated a posteriori (46) followed by standard in-house pipeline of manual inspection and selection of 38,845 micrographs (47) for further processing in cryoSPARC package v4.0.3 (48). Due to the low concentration of particles and the presence of other contaminants in the sample, automatic particle picking appeared challenging, and we developed a CIS-specific approach for efficient picking of the cap and baseplates. In this approach, CIS^Sc^ particles were manually selected as filaments (start-to-end) on 920 micrographs and used for training a crYOLO (49) picking model (Supplementary Fig. 2). The large size of the training set appeared essential for successful picking; we could not get rid of large number of false-positive picks, but there were not so many false-negative picks. Then, CIS^Sc^ particles were picked in crYOLO as filaments, followed by the extraction of the ends of the “filaments” using star_modif.py script (47). Particle coordinates were imported in cryoSPARC, where 211,041 particles were extracted at a large box size of 756 pixels and binned to 64 pixels for initial 2D-classification, which allowed us to separate initial particle sets corresponding to the CIS^Sc^ cap (36,569 particles) and CIS^Sc^ baseplate (43,087 particles) (Supplementary Fig. 2).

Two subsets of this dataset were processed separately in cryoSPARC. For the cap subset, a standard cryoSPARC single-particle processing approach was applied, including gradual iterative 2D-classification rounds, followed by selection of good classes and particle re-extraction at finer sampling. The main challenge was the proper centering of the particles selected on the end of elongated filament-like objects with repeating patterns (sheath-tube module of CIS), hampering the alignments. Ab-initio model generation with applied C6 symmetry was hampered by poor centering of 2D-classes due to elongated nature of the particles (Supplementary Fig. 2). Homo-refinement with this reference was performed followed by particle re-extraction and one more round of ab-initio model generation with applied C6 symmetry (Supplementary Fig. 2). The new 3D-reconstruction appeared reliable and was used for further iterative 3D-refinements (Homo-refinement and NU-refinements) accompanied by 2D-classification and CTF-refinement rounds to exclude particles with wrong Euler angle assignments. The final 3.4 Å resolution map with imposed C6 symmetry resulted from 19,218 particles.

The baseplate subset was processed using the same approach, resulting in a 3D reconstruction with imposed C6 symmetry with a resolution of 3.5Å from 22,980 particles (Supplementary Fig. 2). The tip of the baseplate has a C3 symmetry, therefore to solve that part of the map correctly, we exported that particle stack to Relion4 (50) and took advantage of 2D-classification in Relion to select 18,124 particles, corresponding to the best 2D class averages. 3D-refinement with relaxed symmetry resulted in a 3.8 Å 3D-resolution reconstruction with imposed C3 symmetry.

### Cryo-electron tomography

Intact *Streptomyces* cells were imaged by cryoET as described previously (51). Images were recorded using a Titan Krios 300 kV microscope (Thermo Fisher Scientific) equipped with a Quantum LS imaging filter operated at a 20 eV slit width and with K3 Summit direct electron detectors (Gatan). Tilt series were collected using a bidirectional tilt-scheme from −60 to +60° in 2° increments. Total dose was 130-150 e^-^/Å^2^ and defocus was kept at −8 µm. Tilt series were acquired using SerialEM (52), drift-corrected using alignframes, and reconstructed using IMOD program suite (53). To enhance contrast, tomograms were deconvolved with a Wiener-like filter ‘tom_deconv’ (54).

### Structural modeling of the CIS^Sc^ complex

The map qualities of different modules of CIS^Sc^ (cap and baseplate) allowed for *de novo* structural modeling (Supplementary Fig. 2–3). The atomic models of SCO4243-4248, SCO4252-4253 and SCO4260 were built *de novo* using COOT (55). The resulting models were iteratively refined using RosettaCM (56) and real-space refinement implemented in PHENIX (57). In cases where protein domains could only be partially modeled, side chains were not assigned. Final model validation was done using MolProbity (57) and correlation between models and the corresponding maps were estimated using mtriage (57). To illustrate the complete CIS^Sc^ structure, we generated a composite model by integrating symmetry-related protein subunits into a full model of the CIS^Sc^, based on the consistent CIS^Sc^ length observed both *in situ* and *in vitro* (Fig. 2b).

### Fluorescence microscopy

Fluorescence-based cell viability assays were performed as described previously (18). Briefly, to produce cytoplasmic sfGFP in *Streptomyces*, the coding sequence for sfGFP was introduced downstream of the constitutive promoter *ermE*\* on an integrating plasmid (pIJ10257). The plasmid was introduced by conjugation to *S. coelicolor* strains (SS430, SS575, SS576). An equal level of cytoplasmic sfGFP in the different strains was confirmed by Western blotting (Supplementary Fig. 11). *Streptomyces* strains were grown in 30 ml of TSB liquid culture at 30 °C with shaking at 250 rpm for 48 h. Where appropriate, nisin was added to a final concentration of 1 µg/ml 90 min prior to imaging. One ml aliquots of each culture were vortexed for 1 min to break up mycelial clumps, centrifuged for 5 min at 15,000 x *g*, washed twice with 1xPBS, resuspended in 1 ml of 1xPBS containing 5 µg/ml of the red-fluorescent membrane dye FM5-95 (Invitrogen) and then incubated in the dark at room temperature for 10 min followed by two wash steps with 1x PBS. Washed cell pellets were resuspended in a total volume of 50 µl PBS and 10 µl of each sample was spotted onto 1 % agarose pads and subsequently imaged using the Thunder imager 3D cell culture microscope (Leica). First, tile scan images were acquired using the Las X Navigator plug-in software (Leica Application Suite X, Version 3.7.4.23463), and 100 regions of interest were picked manually. Then Z-stack images were acquired using a HC PL APO 100x objective with the following excitation wavelengths: GFP (475 nm) and TRX (555 nm). Images were processed using LasX software to apply thunder processing and maximum projection and FIJI to create segmentation and quantify the live (sfGFP)/total cells (FM5-95) area ratio (58). Statistical analyses were performed on data from biological triplicates (n=100 images for each experiment) using a one-way ANOVA and Tukey’s post-test in GraphPad Prism 9 (Version 9.3.1).

For imaging the subcellular localization of the CIS adaptor protein in *Streptomyces*, cells (strain JS69 or *S. coelicolor* M145) were grown for 48 h in TSB/YEME liquid medium. Two μl of each strain were spotted on 1 % agarose pads made with water and subsequently imaged.

Images were acquired using a Zeiss Axio Observer Z.1 inverted epifluorescence microscope fitted with a sCMOS camera (Hamamatsu Orca FLASH 4), a Zeiss Colibri 7 LED light source, and a Hamamatsu Orca Flash 4.0v3 sCMOS camera. mCherry fluorescence was detected using an excitation/emission bandwidth of 577–603 nm/614–659 nm. Images were collected using Zen Blue (Zeiss) and analyzed using Fiji (58).

### Cover glass impression of *Streptomyces* spore chains

Spore titers of relevant strains were determined by dilution plating (18). 10^7^ colony-forming units (CFU) of *S. coelicolor* strains (WT, SS539, SS557) were spread onto R2YE agar plates and grown at 30 °C. Sterile glass coverslips were gently applied to the top surface of each bacterial lawn after 48 h, 72 h and 96 h post inoculation. Coverslips were then mounted onto glass microscope slides and imaged using a 40x objective on a Leica Thunder Imager 3D Cell Culture. Images were processed using FIJI (58).

### Actinorhodin production assay

*S. coelicolor* strains (WT, SS539, SS557) were inoculated into 30 ml R2YE liquid media at a final concentration of 1.5 x 10^6^ CFU/ml. Cultures were grown in baffled flasks at 30 °C overnight. Cultures were standardized to an OD_450_ of 0.5 and inoculated in 30 ml of fresh R2YE liquid medium. For the quantification of total actinorhodin production, 480 µl of samples were collected at the indicated time points. 120 µl of 5M KOH was added, samples were vortexed and centrifuged at 5,000 x *g* for 5 min. The weight of each tube was recorded. A Synergy 2 plate reader (BioTek) was used then to measure the absorbance of the supernatant at 640 nm. The absorbance was normalized by the weight of the wet pellet.

### Structure prediction and in silico protein-protein interaction screen

CisA structural models (monomer and oligomers) and the CisA-Cis11 complex were predicted using AlphaFold3 (28) and further processed using UCSF Chimera (59) or ChimeraX-1.7.1 (60).

A pairwise *in silico* screen for possible interactions between CisA and components encoded by the *S. coelicolor cis* gene cluster (*SCO4242-SCO4260*) was conducted using the AlphaFold2 Multimer-based LazyAF pipeline (37,61,62). The confidence metric (ipTM) for the top model from each pairwise interaction was tabulated and visualised using Prism (Version 10.2.2) with ipTM scores >0.7, indicating a possible protein-protein interaction (63).

### Bioinformatic analysis of CisA homologs

To identify CisA homologs, we performed a reciprocal BLAST search in 120 genomes of *Streptomyces* and other *Actinomycete* species that were previously reported to include a type IId eCIS (8,19) using the closely related *cisA* homologue from *Streptomyces albus* J1074 (YP_007743954.1) as a query (72% identical to CisA from *S. coelicolor*) (Supplementary Data 1). Of these 120 genomes, 75 had a reciprocal hit for CisA (Supplementary Data 2). Protein sequences of CisA homologs were aligned using Clustal Omega (64) and visualized with JalView (Version 2.11.3.3).

## Data availability

Representative reconstructed tomograms (EMD-50935, EMD-50939, EMD-50940, EMD-50944, EMD-50948, EMD-50949) and SPA cryoEM maps (EMD-51564, EMD-51565 and EMD-51566) have been deposited in the Electron Microscopy Data Bank. Atomic models (PDB: 9GTP, PDB: 9GTR, PDB: 9GTS) have been deposited in the Protein Data Bank.

## Acknowledgements

We thank ScopeM for instrument access at ETH Zürich. We thank the Functional Genomics Center Zürich for mass spectrometry support. Pilhofer Lab members are acknowledged for discussions. M.P. was supported by the Swiss National Science Foundation (31003A_179255/310030_212592), the European Research Council (679209), and the NOMIS foundation. Work in the lab of S.S. was supported by a Royal Society University Research Fellowship (URF\R1\180075 and URF\R\231009) and BBSRC grant BB/T015349/1 to S.S. and by the BBSRC Institute Strategic Program grant BB/X01097X/1 to the John Innes Centre.

## Author contributions statement

B.C., S.S. and M.P. conceived the project. B.C. optimized the sample preparation for SPA; B.C. and P.A. collected and processed the cryoEM data, reconstructed the cryoEM map; B.C., P.E.H. and J.X. built and refined the structural models; B.C. and P.E.H. conducted and processed cryoET; B.C. conducted automated fluorescent light microscopy, determined sporulation efficiency and actinorhodin production; J.W.S. and S.S. generated plasmids and *Streptomyces* strains; J.W.S. performed membrane topology assays, cellular fractionation experiments and imaged strains expressing fluorescently tagged CisA. S.S. and G.C. performed bioinformatic analyses. B.C., J.W.S., S.S. and M.P. wrote the manuscript.

---

**Supplementary Figure 1:**
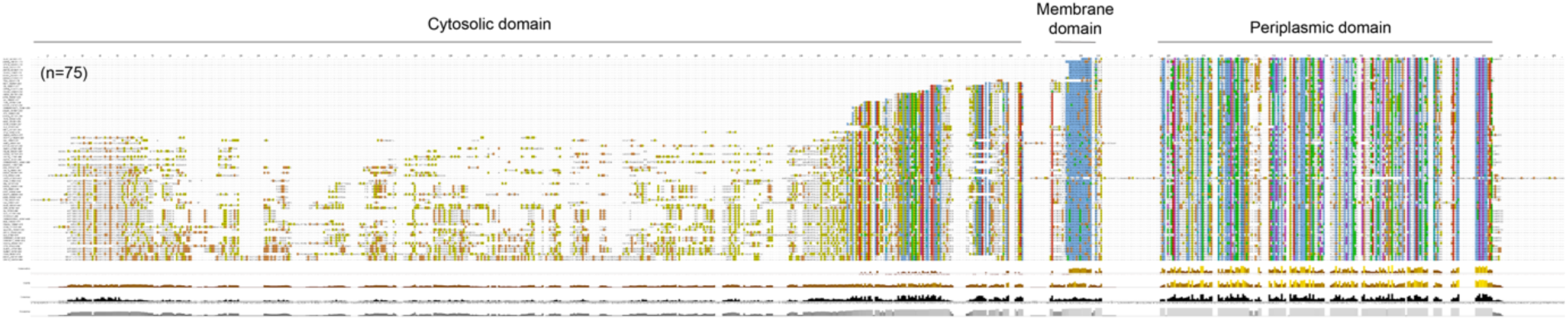
CisA is conserved among CIS-positive *Streptomyces* and Actinomycete species. Protein alignment of CisA homologs identified via reciprocal BLAST search in genomes of *Streptomyces* and actinomycete species previously reported to encode an eCIS class IId region. Protein sequences (n=75) were aligned with Clustal Omega and visualized using JalView. CisA domain structure is indicated on top. Genome accession numbers and BLAST results are listed in Supplementary Data 1 and 2.

**Supplementary Figure 2:**
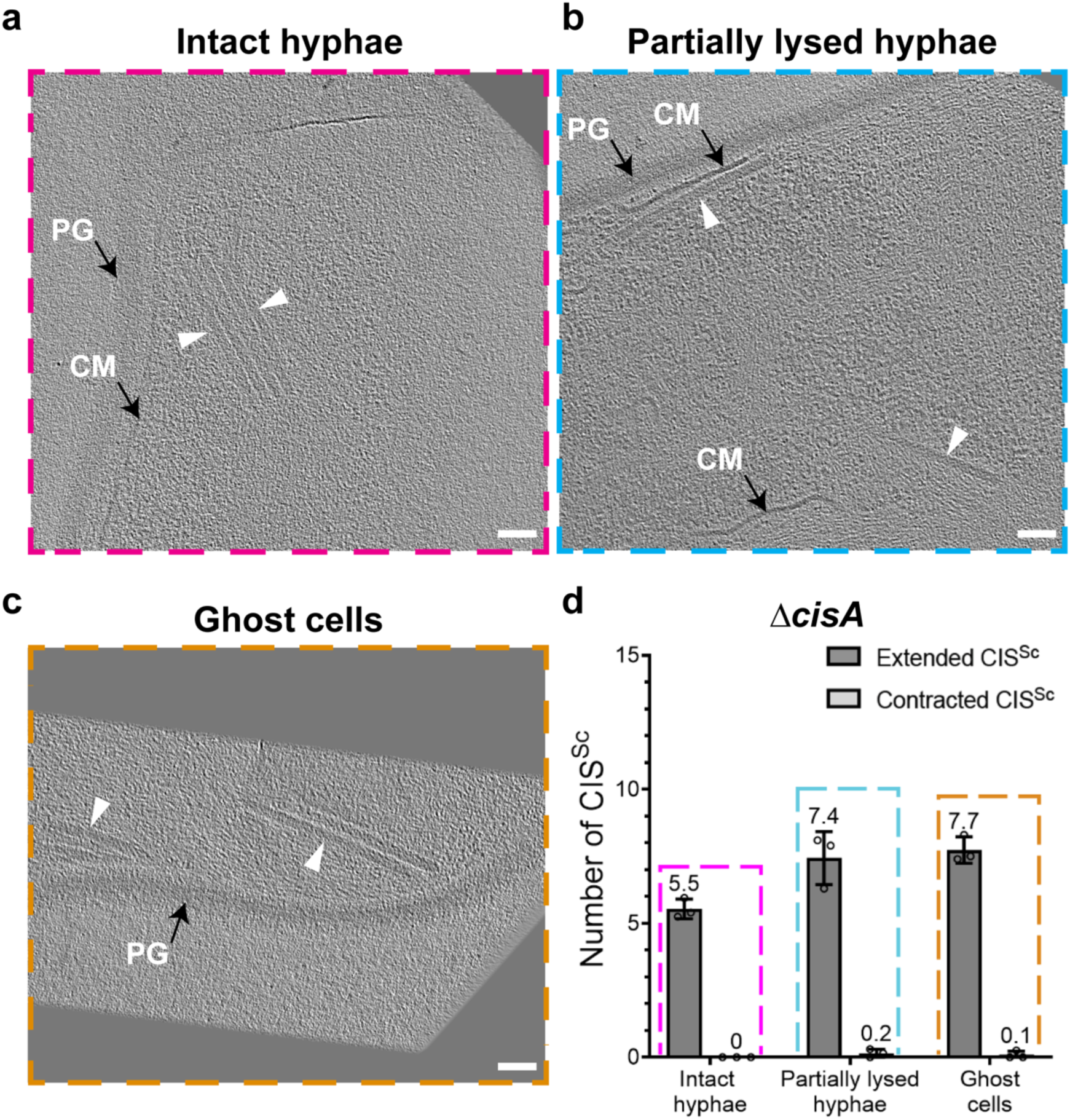
Sheath contraction *in situ* is linked to the presence of CisA. **a-c.** Shown are additional cryo-tomographic slices (thickness 11 nm) of *S. coelicolor ΔcisA* vegetative hyphae, which were observed as ‘Intact hyphae’ (a), ‘Partially lysed hyphae’ (b) and ‘Ghost cells’ (c). Note that all visible CIS^Sc^ particles (white arrowheads) are in the extended conformation. PG, peptidoglycan; CM, cytoplasmic membrane/membranes. Bars, 50 nm. **d.** Shown is a quantification of CIS^Sc^ assemblies per tomogram and their conformations as observed in different classes of Δ*cisA* hyphae. Almost all CIS^Sc^ particles were seen in the extended conformation, indicating that sheath contraction *in situ* correlates with the presence of CisA. Data shows mean values and standard deviations obtained from biological triplicate experiments, with n=30 tomograms for each class of hyphae.

**Supplementary Figure 3:**
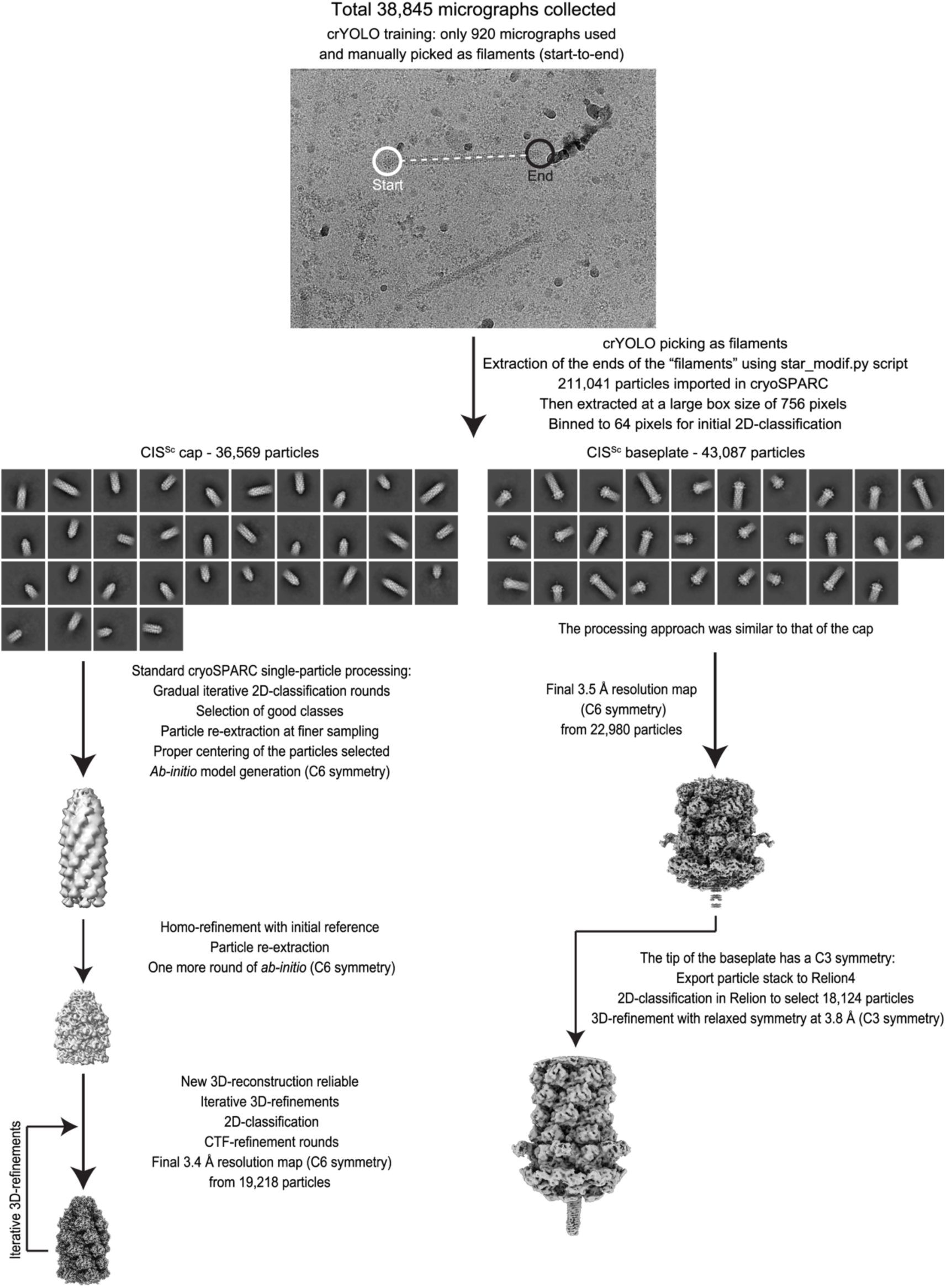
Workflow for the cryoEM structural determination of the extended CIS^Sc^. Flowchart for cryoEM reconstruction of the extended CIS^Sc^ particle. See METHODS and Supplementary Table 1 for details.

**Supplementary Figure 4:**
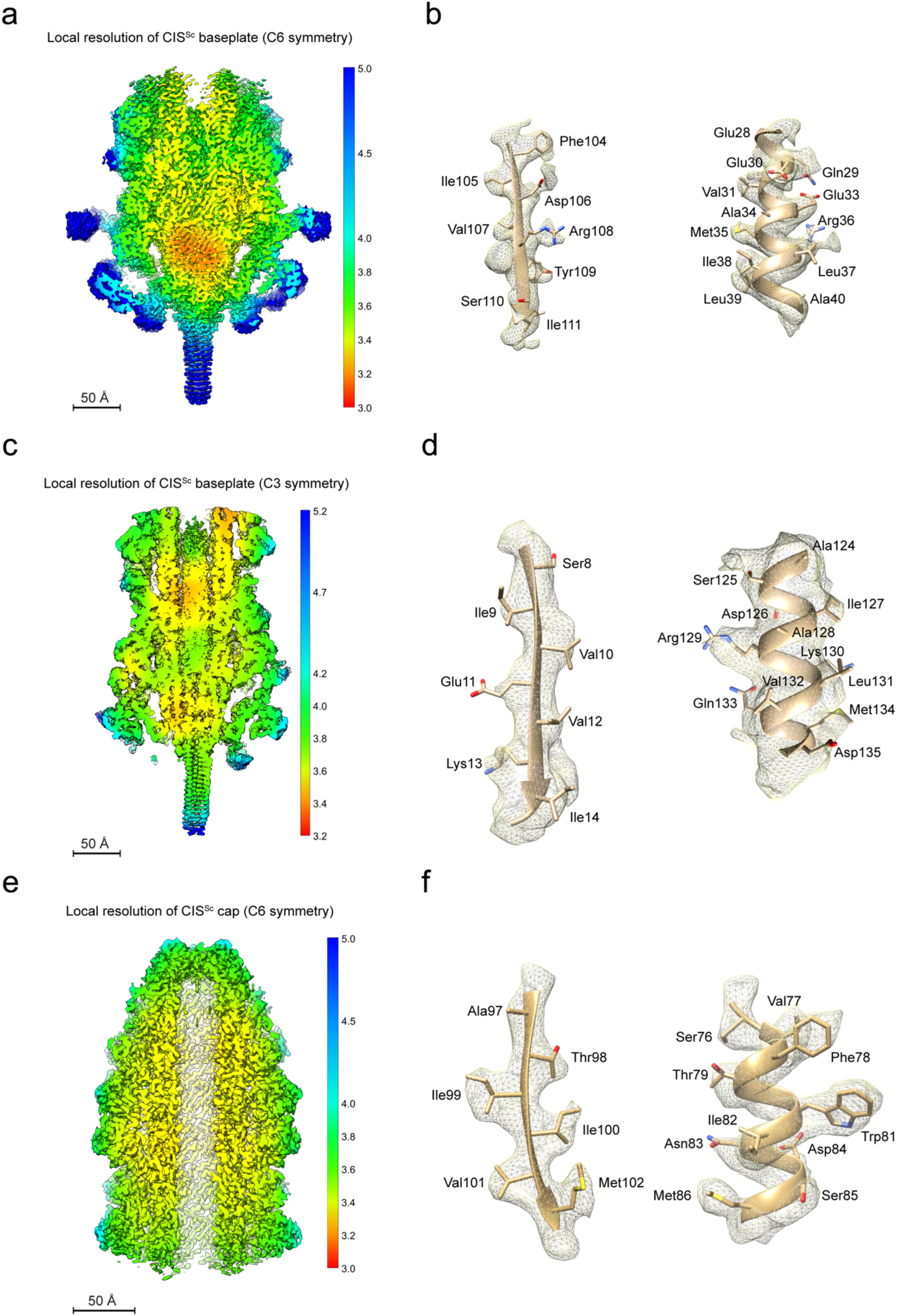
Map qualities of different CIS^Sc^ modules. **a-f.** Local resolution maps of the CIS^Sc^ baseplate with C6 symmetry (a), CIS^Sc^ baseplate with C3 symmetry (c), and the CIS^Sc^ cap with C6 symmetry (e). (d-f) shows regions of the cryoEM density map (mesh) that were superimposed with the atomic models (ribbons and sticks), demonstrating the agreement between the observed and modeled amino acid side chains for one beta-sheet (left) and one alpha helix (right). Shown are examples of the baseplate with C6 symmetry (b), the baseplate with C3 symmetry (d), and the cap with C6 symmetry (f).

**Supplementary Figure 5:**
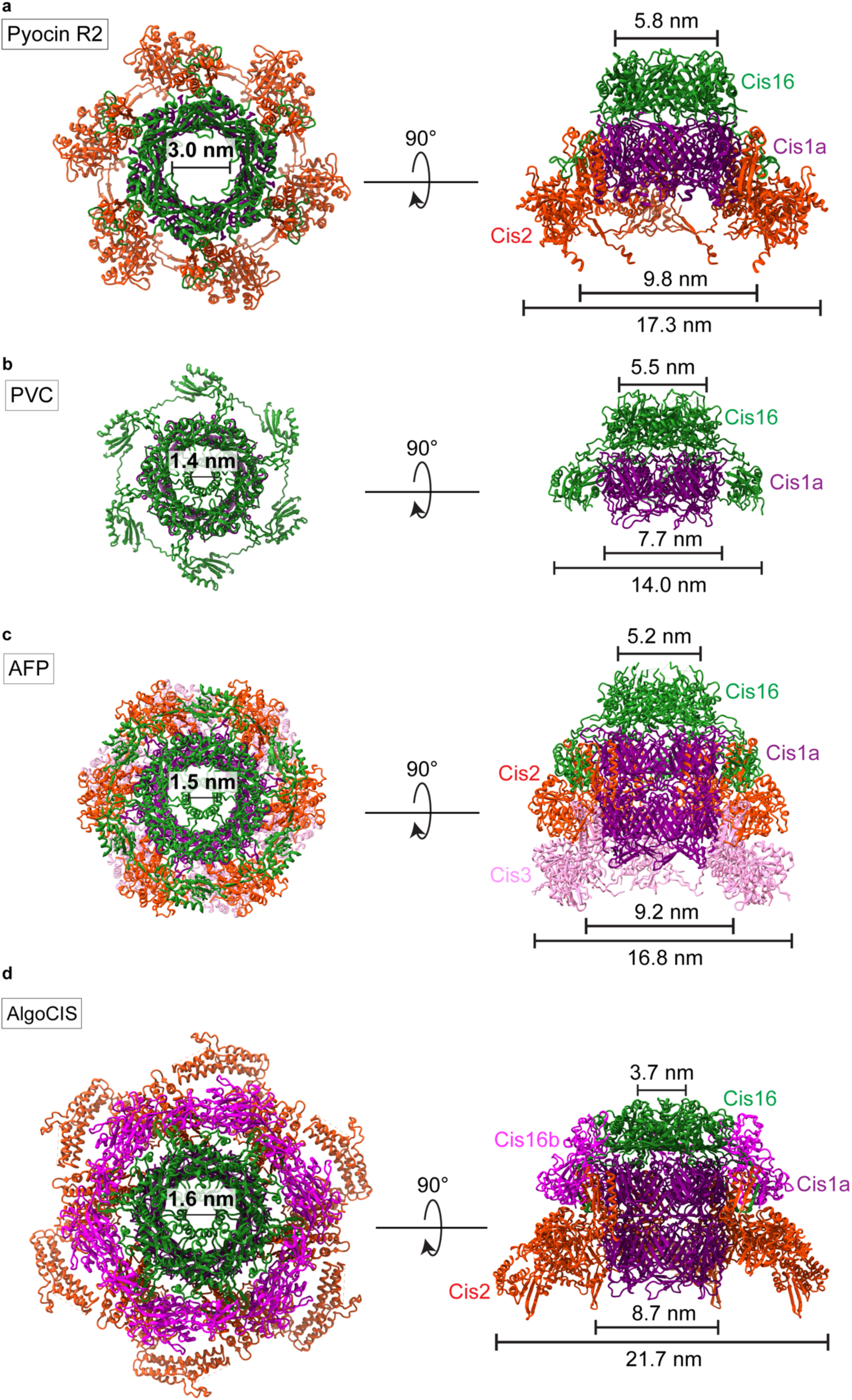
Structures of the cap modules of other CISs. **a-d.** Top views (left) and side views (right) of ribbon representations of the structures of CIS cap modules of Pyocin R2 (a, PDB 6U5B), PVC (b, PDB 6J0N), AFP (**c**, PDB 6RAP) and AlgoCIS (d, PDB 7ADZ).

**Supplementary Figure 6:**
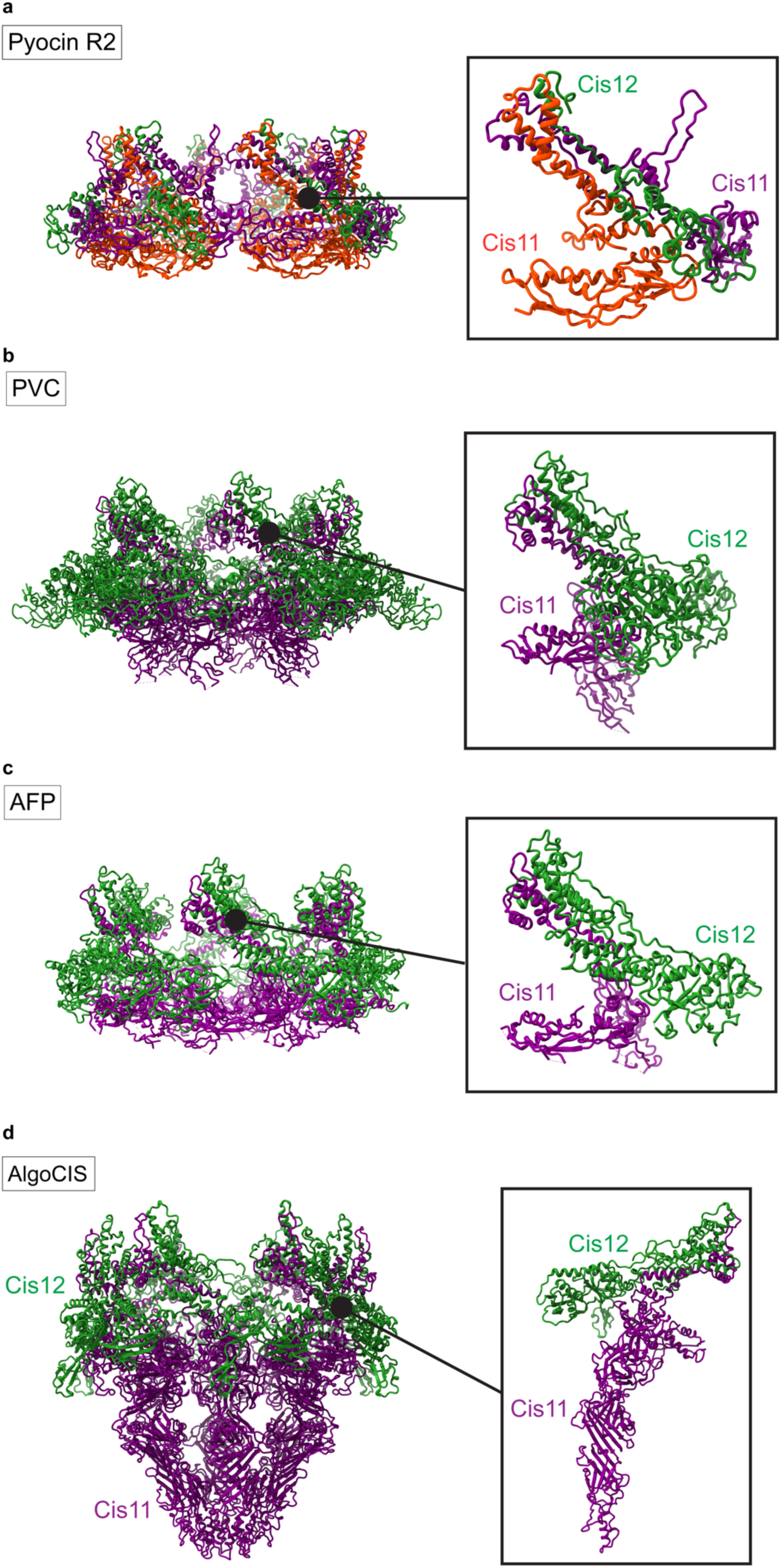
Structures of the baseplate modules of other CISs. **a-d.** Ribbon representations of the full structures (left) and the individual wedge subunit (right) of the CIS baseplate components (Cis11/12) of Pyocin R2 (a, PDB 6U5F), PVC (b, PDB 6J0F), AFP (c, PDB 6RAO) and AlgoCIS (d, PDB 7AEB).

**Supplementary Figure 7:**
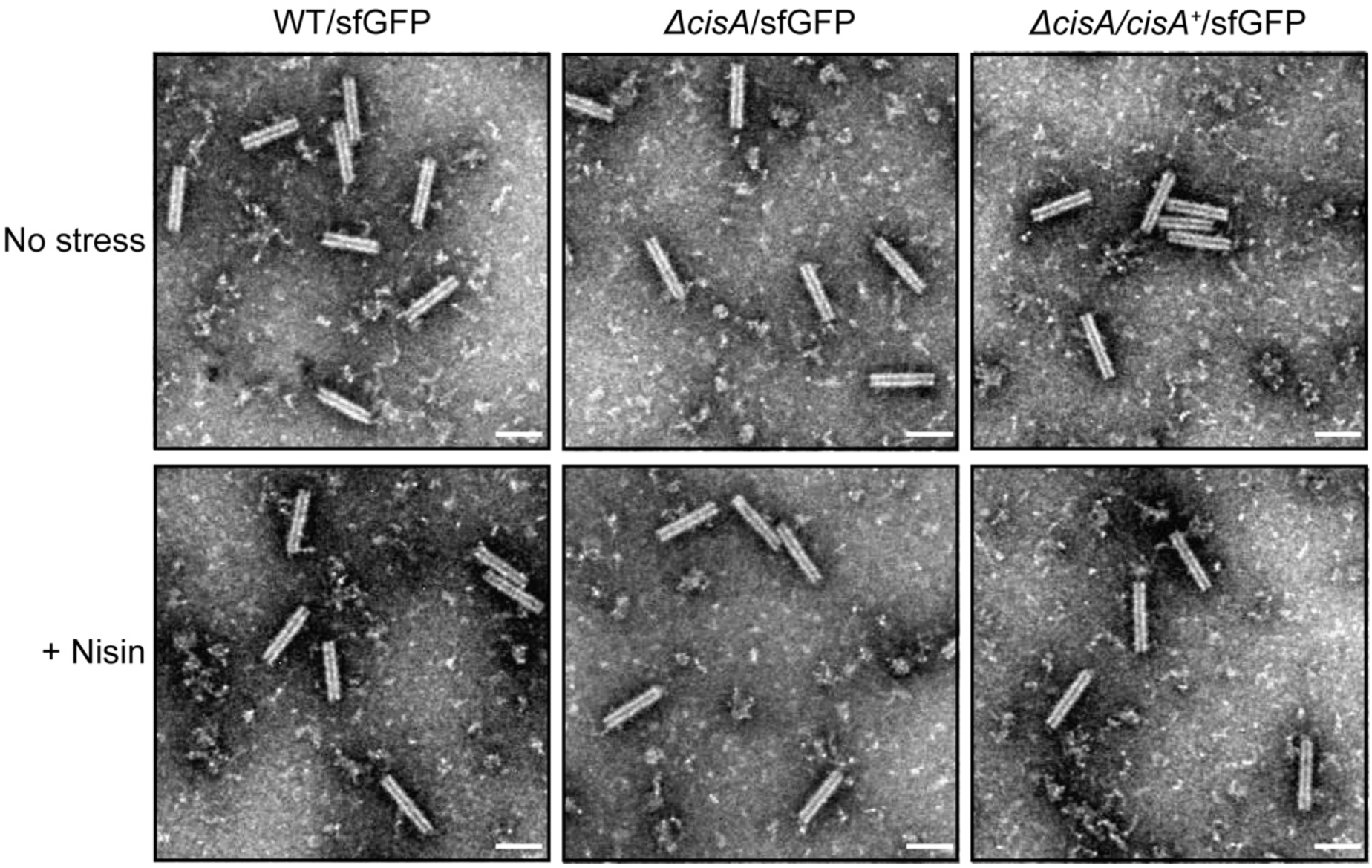
Control experiment showing that the behavior of purified CIS^Sc^ particles is wild-type-like in the strains used in the viability assay. Representative negative-stain electron micrographs of purified CIS^Sc^ particles from the WT/sfGFP, the *ΔcisA*/sfGFP mutant and the *ΔcisA/ΔcisA^+^/*sfGFP complemented mutant exposed to no stress (upper row) or 1 µg/ml nisin (lower row). No differences were observed between the overall structure of CIS^Sc^ particles. These experiments were performed three independent times. Bars, 100 nm.

**Supplementary Figure 8:**
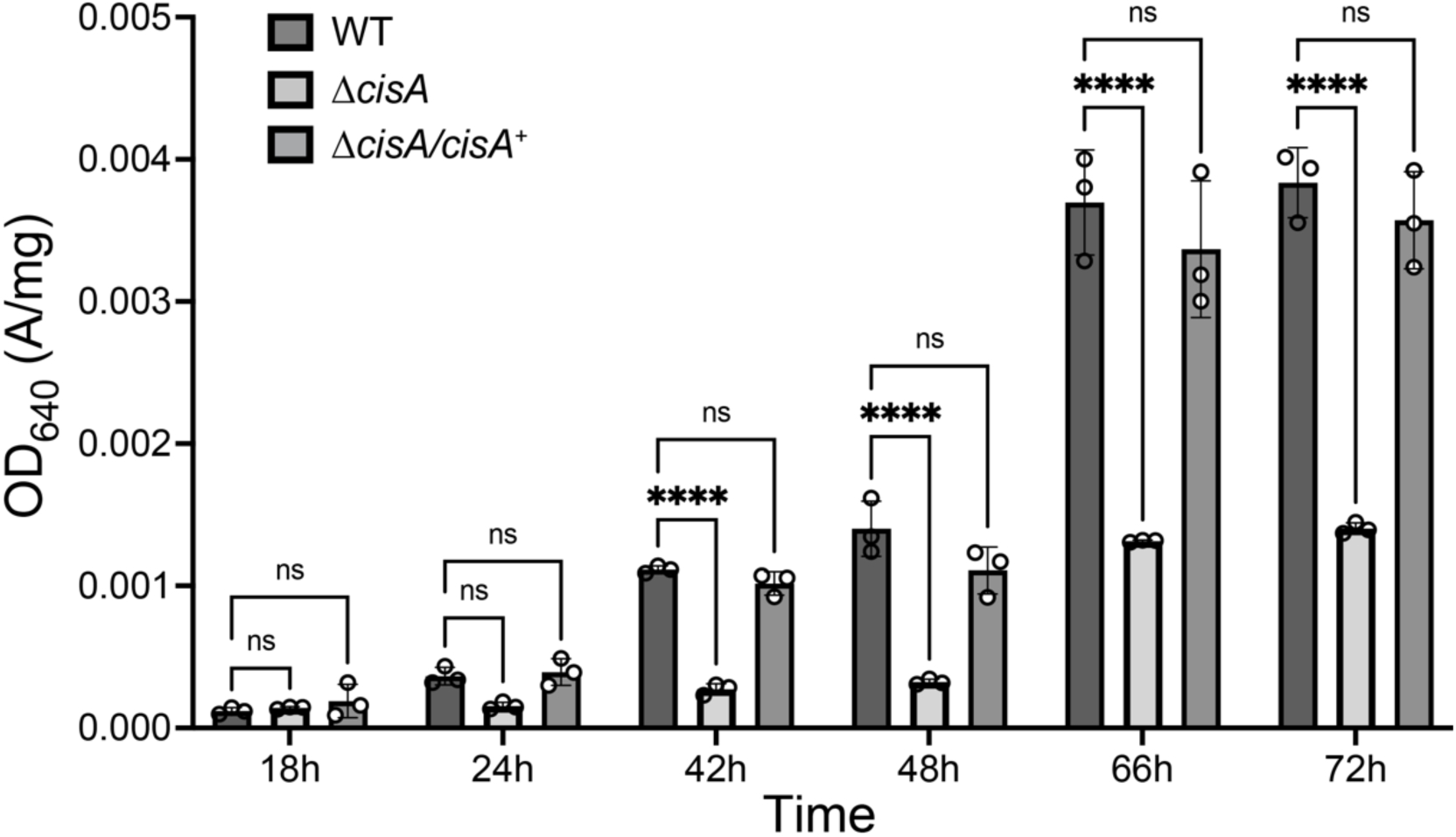
CisA impacts secondary metabolite production. Quantification of total actinorhodin production by the WT, the *ΔcisA* mutant and the complemented *Δcis*A/*ΔcisA^+^* mutant grown in R2YE medium, revealing reduced actinorhodin production in the *ΔcisA* mutant. Samples were taken at the indicated time points. Actinorhodin was extracted and quantified by measuring the optical density OD_640_ of the culture supernatant (65) and normalized to the wet cell pellet weight. Analysis was performed in biological triplicate experiments. The mean values and standard error are shown. ns (not significant) and **** (p < 0.0001) were calculated using one-way ANOVA and Tukey’s post-test.

**Supplementary Figure 9:**
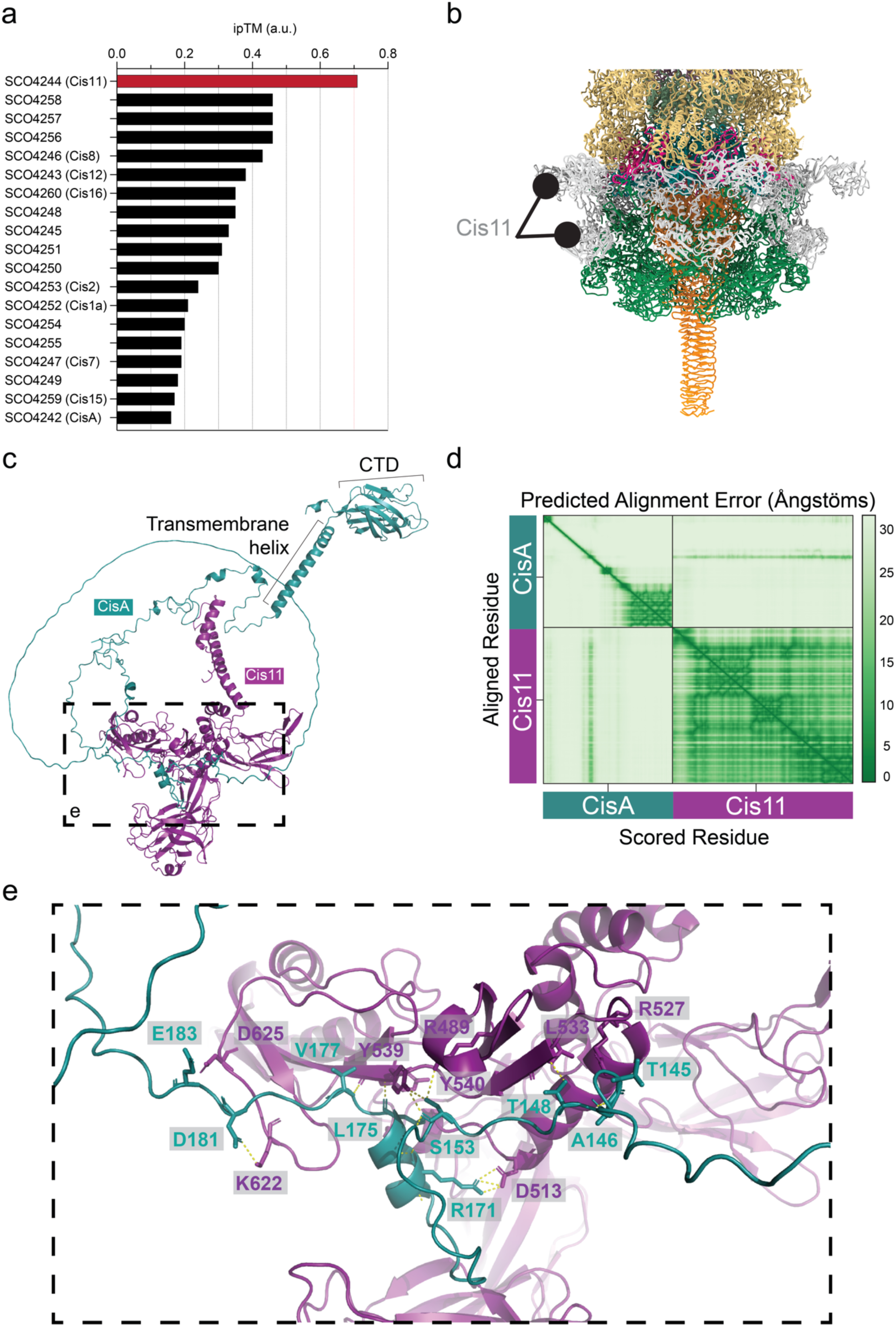
*In silico* protein-protein interaction analyses predict that CisA binds a CIS^Sc^ baseplate component. **a.** An AlphaFold2-Multimer-based protein-protein interaction screen between monomeric CisA and the 19 gene products of the CIS^Sc^ gene cluster suggests an interaction between CisA and the baseplate component Cis11. The likelihood of a protein-protein interaction was ranked based on the interface pTM (ipTM) confidence score, with an ipTM greater than 0.7 indicating a potential protein-protein interaction. **b.** Zoomed-in view of the SPA-cryoEM structure of the CIS^Sc^ baseplate complex (Fig. 2b/d), illustrating the peripheral and surface-exposed position of Cis11. **c.** The Alphafold3 model of the CisA-Cis11 complex. The conserved C-terminal domain (CTD) of CisA is located in the periplasm. **d.** The Alphafold3 predicted aligned error (%) heatmap plot of the concatenated Cis11 and CisA input sequence. **e.** Zoomed-in view of the relevant side chains at the CisA-Cis11 interface.

**Supplementary Figure 10:**
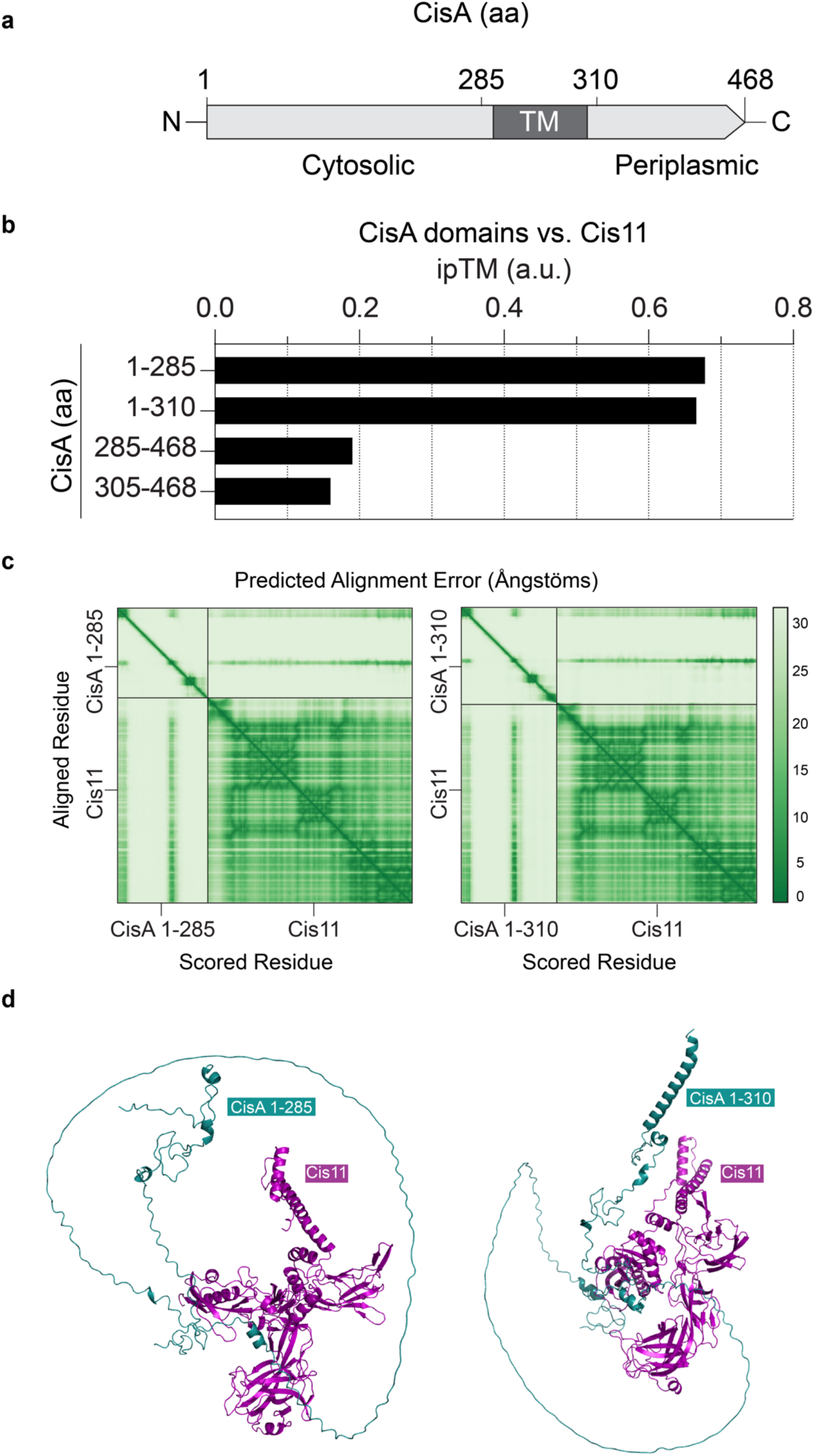
The cytosolic part of CisA is predicted to interact with Cis11. **a.** Schematic showing the CisA domain organization. Relevant amino acid (aa) positions are shown above. TM, transmembrane domain. **b.** AlphaFold2-Multimer based protein-protein interaction screen between monomers of CisA and truncated versions of CisA and Cis11. The likelihood of a protein-protein interaction was ranked based on the interface pTM (ipTM) confidence score. **c/d.** Predicted CisA-Cis11 complexes using truncated versions of CisA (amino acids 1-285 and 1-310) show that the largely unstructured cytosolic portion of CisA is required to interact with Cis11. (c) The Alphafold3 predicted aligned error (%) heatmap plots of the concatenated Cis11 and truncated CisA 1-285 and CisA 1-310 input sequences. (d) The Alphafold3 model of the CisA 1-285-Cis11 (left) and CisA 1-310-Cis11 (right) complexes.

**Supplementary Figure 11:**
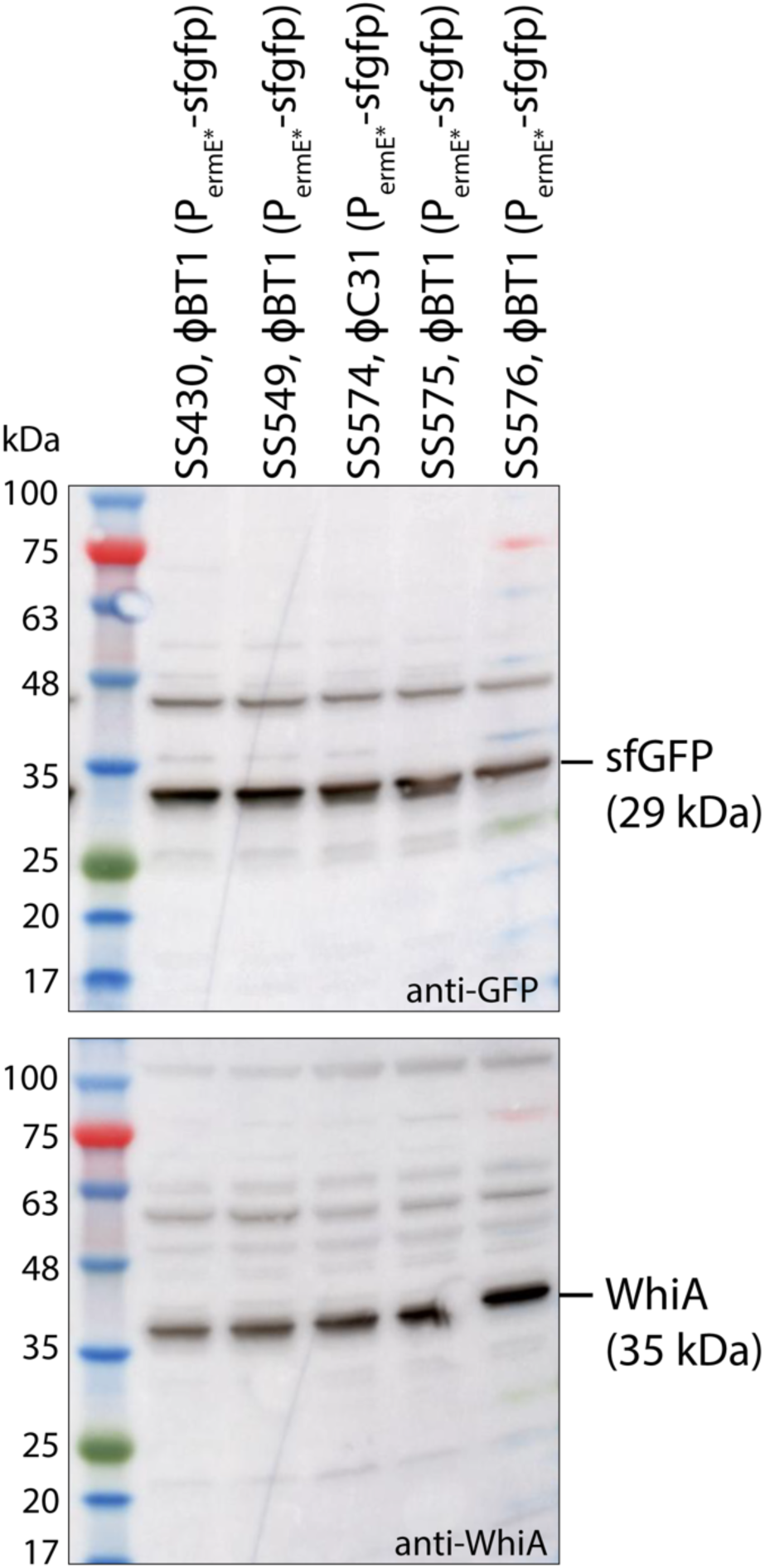
Western blot confirming similar levels of cytosolic sfGFP in strains used for viability assays. Control experiments to verify similar protein levels of sfGFP, which was expressed either from the ϕC31 or the ϕBT1 phage attachment site in the *S. coelicolor* genome. The expression of sfgfp was driven by the constitutive *ermE*\* promoter (*P_ermE*_*). Equal amounts of total protein were loaded and sfGFP levels were detected with an anti-GFP antibody. As a control, the same cell lysates were also probed with an anti-WhiA antibody. Shown are representative results of duplicate experiments.

## References

1. Lin L. The expanding universe of contractile injection systems in bacteria. Curr Opin Microbiol. 2024 Jun 1;79:102465.

2. Brackmann M, Nazarov S, Wang J, Basler M. Using Force to Punch Holes: Mechanics of Contractile Nanomachines. Trends Cell Biol. 2017 Sep;27(9):623–32.

3. Taylor NMI, van Raaij MJ, Leiman PG. Contractile injection systems of bacteriophages and related systems. Mol Microbiol. 2018 Apr;108(1):6–15.

4. Taylor NMI, Prokhorov NS, Guerrero-Ferreira RC, Shneider MM, Browning C, Goldie KN, et al. Structure of the T4 baseplate and its function in triggering sheath contraction. Nature. 2016 May 19;533(7603):346–52.

5. Böck D, Medeiros JM, Tsao HF, Penz T, Weiss GL, Aistleitner K, et al. *In situ* architecture, function, and evolution of a contractile injection system. Science. 2017 Aug 18;357(6352):713–7.

6. Allsopp LP, Bernal P. Killing in the name of: T6SS structure and effector diversity. Microbiology. 2023;169(7):001367.

7. Coulthurst S. The Type VI secretion system: a versatile bacterial weapon. Microbiology (Reading). 2019 May;165(5):503–515.

8. Chen L, Song N, Liu B, Zhang N, Alikhan NF, Zhou Z, et al. Genome-wide Identification and Characterization of a Superfamily of Bacterial Extracellular Contractile Injection Systems. Cell Rep. 2019 Oct 8;29(2):511–521.e2.

9. Geller AM, Pollin I, Zlotkin D, Danov A, Nachmias N, Andreopoulos WB, et al. The extracellular contractile injection system is enriched in environmental microbes and associates with numerous toxins. Nat Commun. 2021 Jun 18;12(1):3743.

10. Basler M, Pilhofer M, Henderson GP, Jensen GJ, Mekalanos JJ. Type VI secretion requires a dynamic contractile phage tail-like structure. Nature. 2012 Feb 26;483(7388):182–6.

11. Durand E, Nguyen VS, Zoued A, Logger L, Péhau-Arnaudet G, Aschtgen MS, et al. Biogenesis and structure of a type VI secretion membrane core complex. Nature. 2015 Jul;523(7562):555–60.

12. Zachs T, Malit JJL, Xu J, Schürch A, Sivabalasarma S, Nußbaum P, et al. Archaeal type six secretion system mediates contact-dependent antagonism. Sci Adv. 2024 Nov 15;10(46):eadp7088.

13. Xu J, Ericson CF, Lien YW, Rutaganira FUN, Eisenstein F, Feldmüller M, et al. Identification and structure of an extracellular contractile injection system from the marine bacterium *Algoriphagus machipongonensis*. Nat Microbiol. 2022 Mar;7(3):397–410.

14. Shikuma NJ, Pilhofer M, Weiss GL, Hadfield MG, Jensen GJ, Newman DK. Marine tubeworm metamorphosis induced by arrays of bacterial phage tail-like structures. Science. 2014 Jan 31;343(6170):529–33.

15. Hurst MRH, Beard SS, Jackson TA, Jones SM. Isolation and characterization of the *Serratia entomophila* antifeeding prophage. FEMS Microbiol Lett. 2007 May;270(1):42– 8.

16. Jiang F, Li N, Wang X, Cheng J, Huang Y, Yang Y, et al. Cryo-EM Structure and Assembly of an Extracellular Contractile Injection System. Cell. 2019 Apr 4;177(2):370–383.e15.

17. Weiss GL, Eisenstein F, Kieninger AK, Xu J, Minas HA, Gerber M, et al. Structure of a thylakoid-anchored contractile injection system in multicellular cyanobacteria. Nat Microbiol. 2022 Mar;7(3):386–96.

18. Casu B, Sallmen JW, Schlimpert S, Pilhofer M. Cytoplasmic contractile injection systems mediate cell death in *Streptomyces*. Nat Microbiol. 2023 Apr;8(4):711–26.

19. Vladimirov M, Zhang RX, Mak S, Nodwell JR, Davidson AR. A contractile injection system is required for developmentally regulated cell death in *Streptomyces coelicolor*. Nat Commun. 2023 Mar 16;14(1):1469.

20. Nagakubo T, Yamamoto T, Asamizu S, Toyofuku M, Nomura N, Onaka H. Phage tail-like nanostructures affect microbial interactions between *Streptomyces* and fungi. Sci Rep. 2021 Oct 11;11:20116.

21. Nagakubo T, Asamizu S, Yamamoto T, Kato M, Nishiyama T, Toyofuku M, et al. Intracellular Phage Tail-Like Nanostructures Affect Susceptibility of *Streptomyces lividans* to Osmotic Stress. mSphere. 2023 Apr 11;8(3):e00114–23.

22. Nagakubo T, Nishiyama T, Yamamoto T, Nomura N, Toyofuku M. Contractile injection systems facilitate sporogenic differentiation of *Streptomyces davawensis* through the action of a phage tapemeasure protein-related effector. Nat Commun. 2024 May 24;15(1):4442.

23. Schlimpert S, Elliot MA. The Best of Both Worlds—*Streptomyces coelicolor* and *Streptomyces venezuelae* as Model Species for Studying Antibiotic Production and Bacterial Multicellular Development. J Bacteriol. 2023 Jun 22;205(7):e00153–23.

24. Flärdh K, Buttner MJ. *Streptomyces* morphogenetics: dissecting differentiation in a filamentous bacterium. Nat Rev Microbiol. 2009 Jan;7(1):36–49.

25. Desfosses A, Venugopal H, Joshi T, Felix J, Jessop M, Jeong H, et al. Atomic structures of an entire contractile injection system in both the extended and contracted states. Nat Microbiol. 2019 Nov 1;4(11):1885–94.

26. Rybakova D, Schramm P, Mitra AK, Hurst MRH. Afp14 is involved in regulating the length of Anti-feeding prophage (Afp). Mol Microbiol. 2015;96(4):815–26.

27. Ge P, Scholl D, Prokhorov NS, Avaylon J, Shneider MM, Browning C, et al. Action of a minimal contractile bactericidal nanomachine. Nature. 2020 Apr;580(7805):658–62.

28. Abramson J, Adler J, Dunger J, Evans R, Green T, Pritzel A, et al. Accurate structure prediction of biomolecular interactions with AlphaFold 3. Nature. 2024 Jun;630(8016):493–500.

29. Alexeyev MF, Winkler HH. Membrane topology of the Rickettsia prowazekii ATP/ADP translocase revealed by novel dual pho-lac reporters 1 1Edited by G. von Heijne. J Mol Biol. 1999 Jan;285(4):1503–13.

30. Ramos-León F, Bush MJ, Sallmen JW, Chandra G, Richardson J, Findlay KC, et al. A conserved cell division protein directly regulates FtsZ dynamics in filamentous and unicellular actinobacteria. eLife. 2021 Mar 17;10:e63387.

31. Bibb MJ, Domonkos A, Chandra G, Buttner MJ. Expression of the chaplin and rodlin hydrophobic sheath proteins in *Streptomyces venezuelae* is controlled by σ(BldN) and a cognate anti-sigma factor, RsbN. Mol Microbiol. 2012 Jun;84(6):1033–49.

32. Malpartida F, Hopwood DA. Molecular cloning of the whole biosynthetic pathway of a *Streptomyces* antibiotic and its expression in a heterologous host. Nature. 1984 May;309(5967):462–4.

33. Rapisarda C, Cherrak Y, Kooger R, Schmidt V, Pellarin R, Logger L, et al. *In situ* and high-resolution cryo-EM structure of a bacterial type VI secretion system membrane complex. EMBO J. 2019 May 15;38(10):e100886.

34. Lien YW, Amendola D, Lee KS, Bartlau N, Xu J, Furusawa G, et al. Mechanism of bacterial predation via ixotrophy. Science. 2024 Oct 18;386(6719):eadp0614.

35. Bongiovanni TR, Latario CJ, Le Cras Y, Trus E, Robitaille S, Swartz K, et al. Assembly of a unique membrane complex in type VI secretion systems of Bacteroidota. Nat Commun. 2024 Jan 10;15(1):429.

36. Ouyang R, Ongenae V, Muok A, Claessen D, Briegel A. Phage fibers and spikes: a nanoscale Swiss army knife for host infection. Curr Opin Microbiol. 2024 Feb 1;77:102429.

37. McLean TC. LazyAF, a pipeline for accessible medium-scale *in silico* prediction of protein-protein interactions. Microbiology (Reading). 2024 Jul;170(7):001473. doi: 10.1099/mic.0.001473.

38. van Kempen M, Kim SS, Tumescheit C, Mirdita M, Lee J, Gilchrist CLM, et al. Fast and accurate protein structure search with Foldseek. Nat Biotechnol. 2024 Feb;42(2):243–6.

39. T. Kieser M. J. Bibb M. J. Buttner K. F. Chater and D. A. Hopwood. Practical Streptomyces Genetics. Norwich: John Innes Foundation; 2000.

40. Gust B, Challis GL, Fowler K, Kieser T, Chater KF. PCR-targeted *Streptomyces* gene replacement identifies a protein domain needed for biosynthesis of the sesquiterpene soil odor geosmin. Proc Natl Acad Sci U S A. 2003 Feb 18;100(4):1541–6.

41. Gust B, Chandra G, Jakimowicz D, Yuqing T, Bruton CJ, Chater KF. Lambda red-mediated genetic manipulation of antibiotic-producing *Streptomyces*. Adv Appl Microbiol. 2004;54:107–28.

42. Ohi M, Li Y, Cheng Y, Walz T. Negative Staining and Image Classification - Powerful Tools in Modern Electron Microscopy. Biol Proced Online. 2004;6:23–34.

43. Bush MJ, Bibb MJ, Chandra G, Findlay KC, Buttner MJ. Genes Required for Aerial Growth, Cell Division, and Chromosome Segregation Are Targets of WhiA before Sporulation in *Streptomyces venezuelae*. mBio. 2013 Sep 24;4(5):e00684–13.

44. Gallego JJ, Severi E, Chandra G, Palmer T. Identification of novel tail-anchored membrane proteins integrated by the bacterial twin--arginine translocase. Microbiology (Reading). 2024 Feb;170(2):001431.

45. Zheng SQ, Palovcak E, Armache JP, Verba KA, Cheng Y, Agard DA. MotionCor2: anisotropic correction of beam-induced motion for improved cryo-electron microscopy. Nat Methods. 2017 Apr;14(4):331–2.

46. Afanasyev P, Ravelli RBG, Matadeen R, De Carlo S, van Duinen G, Alewijnse B, et al. A posteriori correction of camera characteristics from large image data sets. Sci Rep. 2015 Jun 11;5(1):10317.

47. 47. P. Afanasyev. CryoEM tools [Internet]. 2023. Available from: https://github.com/afanasyevp/cryoem_tools

48. Punjani A, Rubinstein JL, Fleet DJ, Brubaker MA. cryoSPARC: algorithms for rapid unsupervised cryo-EM structure determination. Nat Methods. 2017 Mar 1;14(3):290–6.

49. Wagner T, Merino F, Stabrin M, Moriya T, Antoni C, Apelbaum A, et al. SPHIRE-crYOLO is a fast and accurate fully automated particle picker for cryo-EM. Commun Biol. 2019 Jun 19;2(1):1–13.

50. Scheres SHW. RELION: implementation of a Bayesian approach to cryo-EM structure determination. J Struct Biol. 2012 Dec;180(3):519–30.

51. Weiss GL, Medeiros JM, Pilhofer M. In Situ Imaging of Bacterial Secretion Systems by Electron Cryotomography. Methods Mol Biol Clifton NJ. 2017;1615:353–75.

52. Mastronarde DN. Automated electron microscope tomography using robust prediction of specimen movements. J Struct Biol. 2005 Oct;152(1):36–51.

53. Kremer JR, Mastronarde DN, McIntosh JR. Computer visualization of three-dimensional image data using IMOD. J Struct Biol. 1996 Feb;116(1):71–6.

54. Tegunov D, Cramer P. Real-time cryo-electron microscopy data preprocessing with Warp. Nat Methods. 2019 Nov;16(11):1146–52.

55. Emsley P, Lohkamp B, Scott WG, Cowtan K. Features and development of Coot. Acta Crystallogr D Biol Crystallogr. 2010 Apr;66(Pt 4):486–501.

56. Song Y, DiMaio F, Wang RYR, Kim D, Miles C, Brunette T, et al. High-resolution comparative modeling with RosettaCM. Struct Lond Engl 1993. 2013 Oct 8;21(10):1735–42.

57. Adams PD, Afonine PV, Bunkóczi G, Chen VB, Davis IW, Echols N, et al. PHENIX: a comprehensive Python-based system for macromolecular structure solution. Acta Crystallogr D Biol Crystallogr. 2010 Feb;66(Pt 2):213–21.

58. Schindelin J, Arganda-Carreras I, Frise E, Kaynig V, Longair M, Pietzsch T, et al. Fiji - an Open Source platform for biological image analysis. Nat Methods. 2012 Jun 28;9(7):10.1038/nmeth.2019.

59. Ef P, Td G, Cc H, Gs C, Dm G, Ec M, et al. UCSF Chimera - a visualization system for exploratory research and analysis. J Comput Chem. 2004 Oct 2022 May 17];25(13).

60. Goddard TD, Huang CC, Meng EC, Pettersen EF, Couch GS, Morris JH, et al. UCSF ChimeraX: Meeting modern challenges in visualization and analysis. Protein Sci Publ Protein Soc. 2018 Jan;27(1):14–25.

61. Jumper J, Evans R, Pritzel A, Green T, Figurnov M, Ronneberger O, et al. Highly accurate protein structure prediction with AlphaFold. Nature. 2021 Aug;596(7873):583–9.

62. Mirdita M, Schütze K, Moriwaki Y, Heo L, Ovchinnikov S, Steinegger M. ColabFold: making protein folding accessible to all. Nat Methods. 2022 Jun;19(6):679–82.

63. O’Reilly FJ, Graziadei A, Forbrig C, Bremenkamp R, Charles K, Lenz S, et al. Protein complexes in cells by AI-assisted structural proteomics. Mol Syst Biol. 2023 Apr 12;19(4):e11544.

64. Madeira F, Madhusoodanan N, Lee J, Eusebi A, Niewielska A, Tivey ARN, et al. The EMBL-EBI Job Dispatcher sequence analysis tools framework in 2024. Nucleic Acids Res. 2024 Jul 5;52(W1):W521–5.

65. Zhang Y, Wang L, Zhang S, Yang H, Tan H. hmgA, transcriptionally activated by HpdA, influences the biosynthesis of actinorhodin in *Streptomyces coelicolor*. FEMS Microbiol Lett. 2008 Mar;280(2):219–25.

