## Supplementary Tables for "Function and firing of the *Streptomyces coelicolor* contractile injection system requires the membrane protein CisA"

**Supplementary Table 1.** Cryo-EM data statistical analysis

| | Cap | Baseplate_C6 (overall) | Baseplate_C3 |
|---|---|---|---|
| **Data collection and processing** | | | |
| Nominal magnification | | 81,000 | |
| Voltage (kV) | | 300 | |
| Electron exposure ($e^-$/Å) | | ~60 (K3 camera) | |
| Defocus range (μm) | | 1.0 - 3.0 | |
| Pixel size (Å/pixel) | | 1.065 | |
| Symmetry imposed | C6 | C6 | C3 |
| Initial particles (No.) | 36,569 | 43,087 | 36,569 |
| Final particles (No.) | 19,218 | 22,920 | 18,124 |
| Map resolution (Å) | 3.4 | 3.5 | 3.8 |
| FSC threshold | | 0.143 | |
| **Refinement** | | | |
| Model composition | | | |
| Atoms | 33006 (Hydrogens: 243) | 141138 (Hydrogens: 0) | 14343 (Hydrogens: 0) |
| Protein residues | 4266 | 18408 | 1923 |
| Chains | 18 | 60 | 3 |
| R.M.S deviations | | | |
| Bond length (Å) | 0.003 | 0.007 | 0.005 |
| Bond angles (˚) | 0.558 | 0.839 | 0.768 |
| Validation | | | |
| MolProbity score | 1.51 | 2.22 | 1.91 |
| Clashscore | 5.77 | 14.01 | 7.86 |
| Rotamer outlier (%) | 0.03 | 0.04 | 0 |
| Ramachandran plot | | | |
| Favored (%) | 96.81 | 89.51 | 92.18 |
| Allowed (%) | 3.19 | 10.46 | 7.82 |
| Outlier (%) | 0 | 0.44 | 0 |
| Masked CC | 0.77 | 0.79 | 0.77 |

**Supplementary Table 2.** Proteins detected by mass spectrometry in samples of purified $CIS^{Sc}$ after crude sheath preparation from a *S. coelicolor* non-contractile $CIS^{Sc}$ mutant without exogeneous stress and following nisin stress.

| Protein ID | CIS ID | *S. coelicolor* CIS-N5 | |
|---|---|---|---|
| | | Without exogeneous stress | Upon nisin stress |
| SCO4242 | CisA | - | 5% coverage / 2 total unique peptide |
| SCO4243 | Cis12 | 6% coverage / 1 total unique peptide | 5% coverage / 1 total unique peptide |
| SCO4244 | Cis11 | 13% coverage / 6 total unique peptide | 4% coverage / 2 total unique peptide |
| SCO4245 | Cis9 | 21% coverage / 2 total unique peptide | 20% coverage / 2 total unique peptide |
| SCO4246 | Cis8 | 18% coverage / 10 total unique peptide | 13% coverage / 7 total unique peptide |
| SCO4247 | Cis7 | 5% coverage / 1 total unique peptide | 4% coverage / 1 total unique peptide |
| SCO4248 | Cis1b | 22% coverage / 2 total unique peptide | 20% coverage / 2 total unique peptide |
| SCO4249 | - | 5% coverage / 1 total unique peptide | 4% coverage / 1 total unique peptide |
| SCO4252 | Cis1a | 26% coverage / 4 total unique peptide | 19% coverage / 3 total unique peptide |
| SCO4253 | Cis2 | 23% coverage / 9 total unique peptide | 30% coverage / 14 total unique peptide |
| SCO4254 | - | 10% coverage / 6 total unique peptide | 10% coverage / 6 total unique peptide |
| SCO4256 | - | 17% coverage / 7 total unique peptide | 22% coverage / 9 total unique peptide |
| SCO4257 | - | - | - |
| SCO4258 | - | - | - |
| SCO4259 | Cis15 | - | - |
| SCO4260 | Cis16 | 20% coverage / 4 total unique peptide | 17% coverage / 3 total unique peptide |

**Supplementary Table 3:** Strains and plasmids used in this study

| Strain | Description | Construction | Source |
|---|---|---|---|
| ***Escherichia coli* strains** | | | |
| TOP10 | *F⁻ mcrA Δ(mrr-hsdRMS-mcrBC) Φ80lacZΔM15 ΔlacX74 recA1 araD139 Δ(ara leu) 7697 galU galK rpsL (StrR ) endA1 nupG* | Cloning | Lab strain |
| ET12567/pUZ8002 | *F⁻ dam13::Tn9 dcm6 hsdM hsdR recF143:: Tn10 galK2 galT22 ara-14 lacY1 xyl-5 leuB6 thi-1 tonA31 rpsL hisG4 tsx-78 mtl-1 glnV44* | ET12567 with helper plasmid pUZ8002 | (1) |
| BW25113/pIJ790 | *Δ(araD-araB)567 ΔlacZ4787(::rrnB-4) lacIp-4000(lacIQ), λ−rpoS369(Am) rph-1 Δ(rhaD-rhaB)568 hsdR514* | BW25113 containing λ RED recombination plasmid pIJ790 | (2) |
| NEB5-alpha | *fhuA2Δ(argF-lacZ)U169 phoA glnV44 Φ80 Δ(lacZ)M15 gyrA96 recA1 relA1 endA1 thi-1 hsdR17* | Cloning | NEB |
| Rosetta (DE3) | *$F^-$ ompT hsdSB(rB$^-$ mB-) gal dcm (DE3) pRARE (Cam$^R$)* | Host strain for protein overexpression | Merck |
| TG1 | *supE thi-1 Δ(lac-proAB) Δ(mcrB-hsdSM)5 (rK-mK) / F´ traD36 proAB lacIqZΔM15* | Topology Experiment | Lab Strain |
| ***Streptomyces coelicolor* strains** | | | |
| *S. coelicolor* M145 | Wild type (SCP1-, SCP2-) | | (3) |
| SS387 | Δ*SCO4253-51::apr, apr$^R$* | | (4) |

| | | | |
|---|---|---|---|
| SS393 | *ΔSCO4253-51::apr attB ϕBT1 (SCO4253-N5-SCO4252-51), hyg$^R$* | | (4) |
| SS430 | WT *ϕBT1 (PermE*-sfgfp)* | | (4) |
| SS539 | *ΔSCO4242::apr, apr$^R$* | chromosomal *SCO4242 (cisA)* locus deleted using pSS684 | This study |
| SS549 | *ΔSCO4242::apr ϕBT1 (PermE*-sfgfp)* | pSS150 integrated at ΦBT1 attachment site of SS539 | This study |
| SS557 | *ΔSCO4242 ϕBT1(PermE*-SCO4242-3xFLAG) hyg$^R$, apr$^R$* | pSS735 integrated at ϕBT1 attachment site of SS539 | This study |
| SS574 | WT *ϕC31 (PermE*-sfgfp), apr$^R$, thio$^R$* | pSS758 integrated at ϕC31 attachment site of WT | This study |
| SS575 | *ΔSCO4242::apr ϕC31 (PermE*-sfGFP) apr$^R$, thio$^R$* | pSS758 integrated at ϕC31 attachment site of SS539 | This study |
| SS576 | *ΔSCO4242:apr ϕBT1(PermE*-SCO4242-3xFLAG) ϕC31 (PermE*-sfGFP) apr$^R$, thio$^R$, hyg$^R$* | pSS758 integrated at ϕC31 attachment site of SS557 | This study |
| JS65 | *ΔSCO4242:apr ϕBT1(PermE*-) hyg$^R$, apr$^R$* | pIJ10257 integrated at ϕBT1 attachment site of SS387 | This Study |
| JS69 | *ΔSCO4242:apr ϕBT1(PermE*-SCO4242-mcherry) hyg$^R$, apr$^R$* | pSS705 integrated at ϕBT1 attachment site of SS387 | This study |
| **Cosmid and Plasmids** | | | |
| StD8A | Cosmid vector containing coding sequence for *SCO4242* km$^R$, carb$^R$ | | http://strepdb.streptomyces.org.uk |
| pIJ773 | pBluescript KS (+) containing the apramycin resistance gene *apr* and *oriT* of plasmid RP4, | | (5) |

| | | | |
|---|---|---|---|
| | flanked by FRT sites (Apr$^R$). Used as template for the amplification of the *apr-oriT* cassette for 'REDIRECT' PCR targeting, apr$^R$ | | |
| pIJ10257 | Cloning vector for the conjugal transfer of DNA (under control of the *ermE** constitutive promoter). Integrates at the *ΦBT1* attachment site, hyg$^R$ | | (6) |
| pET21b | Plasmid for overexpression of proteins in *E. coli*, *carb$^R$* | | Lab stock |
| pKF351 | Derivative of pKF280, integrates at the *ΦC31* attachment site, apr$^R$ | | (7) |
| pSS150 | pIJ10257 carrying *PermE*-sfgfp*, hyg$^R$ | | (4) |
| pSS684 | Mutated cosmid StD8A for REDIRECT containing *ΔSCO4242::apr, km$^R$, carb$^R$, apr$^R$* | The *SCO4242* coding sequence on the cosmid vector StD8A was replaced by an *oriT*-containing apramycin resistance cassette, which amplified from pIJ773 using primer 1676/1677 | This study |
| pSS88 | pIJ10257 carrying *mcherry*, *hyg$^R$* | *Streptomyces* Codon-optimised *mcherry* was inserted between the NdeI-XhoI site of pIJ10257. | This study |
| pSS705 | *PermE*-SCO4242-mcherry, hyg$^R$* | *SCO4242* coding sequence amplified with primer 1731/1732 and assembled with pSS88 cut with NdeI-XhoI. | This study |
| pSS734 | *PermE*-SCO4242- TEV-sfgfp, hyg$^R$* | A "GGS-TEV" amino acid sequence was added upstream of *sfgfp* from pSS150 by PCR using | This study |

| | | | |
|---|---|---|---|
| | | primer 1800/1801. The PCR product was cut with XhoI-AvrI and inserted into pSS705 cut with the same enzymes. | |
| pSS735 | *PermE*-SCO4242-3xFLAG, hyg*$^R$ | *PermE*-SCO4242* fragment was isolated from pSS705 by restriction digestion with KpnI-XhoI and ligated with pIJ10257 cut with KpnI-XhoI. | This study |
| pSS758 | pKF351 carrying *PermE*-sfgfp, apr*$^R$*, thio*$^R$ | *sfgfp* amplified from pSS150 with primer 1851/1852 and assembled into pKF351 cut with BamHI-KpnI. | This study |
| **Plasmid used to test membrane topology** | | | |
| pKTop | Plasmid encoding a dual *pho-lac* reporter, km$^R$ | | (8) |
| pFRL1 | Plasmid encoding the known *Streptomyces* cytoplasmic protein, SepH, in the dual *pho-lac* reporter, *km*$^R$ | | Felix Ramos-Leon (unpublished) |
| pFRL5 | Plasmid encoding the *Streptomyces* membrane protein, RsbN, in the dual *pho-lac* reporter, *km*$^R$ | | Felix Ramos-Leon (unpublished) |
| pJUK131 | pKTop carrying *SCO4242, km*$^R$ | Codon optimized *SCO4242* amplified with primers JS111/JS112 from pJUK123 digested and inserted into *HinDIII*/*Kpn*I digested pKTop | This study |
| pJUK132 | pKTop carrying *SCO4242* (1-310aa), *km*$^R$ | Codon optimized *SCO4242* containing only codons for aa 1-310 amplified with primers JS111/JS113 from pJUK123 digested and inserted into *HinDIII*/*Kpn*I digested pKTop | This study |
| pJUK145 | pKTop carrying *SCO4242* (1-285aa), *km*$^R$ | Codon optimized *SCO4242* containing only codons for aa 1-285 amplified with primers JS111/JS114 from pJUK123 digested and inserted into HinDIII/KpnI digested pKTop | This study |

| | | | |
|---|---|---|---|
| pJUK146 | pKTop carrying *SCO4242* (Δ295-305aa), *km*$^R$ | Codon optimized *SCO4242* omitting codons for aa 295-310 amplified with primers JS111/JS112 from pJUK133 digested and inserted into HinDIII/KpnI digested pKTop | This study |

**Supplementary Table 4**: Oligonucleotides used in this study.

| Name | Sequence (5'→3') |
|---|---|
| 1676 | GGAGAGATGACCACGCAGAACTGCGCCGAGTGCGGAACCATTCCGGGGATCCGTCGACC |
| 1677 | CGGCCCGGGTCAGGAGGAGCGGTTGGCGCTGGACGGGCCTGTAGGCTGGAGCTGCTTC |
| 1691 | ATATTCTAGAGATGACCACGCAGAACTGCGCC |
| 1692 | ATATGGTACCTCAGGAGGAGCGGTTGGCGC |
| 1693 | ATATAAGCTTGATGCCCCTGCCCTCTCCCAAC |
| 1694 | ATATGGTACCCGCGCGCTGTCCCCGATCACG |
| 1731 | TGGTAGGATCGTCTAGAACAGGAGGCCCCATATGACCACGCAGAACTGCGCC |
| 1732 | ATGTTGTCCTCCTCGCCCTTGGAGACCATCTCGAGGGAGGAGCGGTTGGCGCT |
| 1767 | AGCTGTTTCCTGTGTGAAATTGTTATCC |
| 1800 | ATATACTCGAGGGTGGCTCCGAAAACCTGTACTTCCAATCCATGTCCAAGGGCGAGGAGC |
| 1801 | AATTAACCTAGGTCACTTGTACAGCTCGTCCATGCC |
| 1851 | AAGTCGTGCTGCTTCATGTGGTCGGGGTACCCGACCCGAGCACGCG |
| 1852 | ATTACTGGACCGGATGAATTCACTTGGATCCGTTAATTAATCACTCGAGCTCCGGGCCCG |
| 1930 | CCAAGCTTGCATGCCTGCAGG |
| 1931 | TCACACAGGAAACAGCTATGACCACGCAGAACTGCGCC |
| 1932 | GGCATGCAAGCTTGGGATGCGGTCGAAGACGCGGC |
| 1933 | CCAAGCTTGCATGCCTGCAGG |
| 1934 | GGCATGCAAGCTTGGGATGCGGTCGAAGACGCGGC |
| JS105 | CCTCCGCGAATCGGTCGTCTCTCGAGGGTGGCTCCGAAAACCT |
| JS106 | GCGCAATTTTGGGTCGTCATATGTATATCTCCTTCTTAAAGTTAAACAAAATTATTTCTAGAGGGGAA |
| JS107 | TTTAAGAAGGAGATATACATATGACGACCCAAAATTGCGCCGAATGT |
| JS108 | TTTTCGGAGCCACCCTCGAGAGACGACCGATT |
| JS111 | CATTAGAAGCTTTATGACGACCCAAAATTGCGCCG |
| JS112 | CATTAGGGTACCGACGACCGATTCGCGGAGG |
| JS113 | CATTAGGGTACCTGAACCCCGTCGGGAATATA |
| JS114 | CATTAGGGTACCCCGATTCGATCGAAAACACGACGC |

**Supplementary Data 1**: List of all genomes used for reciprocal BLAST search to identify CisA homologs.

**Supplementary Data 2:** BLAST search results for homologues of CisA homologs. Streptomyces albus J1074 (YP_007743954.1) was used as a query. YP_007743954.1 is closely related to SCO4242/CisA (72% identical).

**References:**

1. Paget MSB, Chamberlin L, Atrih A, Foster SJ, Buttner MJ. Evidence that the extracytoplasmic function sigma Factor $\sigma^E$ is required for normal cell wall structure in *Streptomyces coelicolor* A3(2). J Bacteriol. 1999 Jan;181(1):204–11.

2. Datsenko KA, Wanner BL. One-step inactivation of chromosomal genes in *Escherichia coli* K-12 using PCR products. Proc Natl Acad Sci. 2000 Jun 6;97(12):6640–5.

3. T. Kieser M. J. Bibb M. J. Buttner K. F. Chater and D. A. Hopwood. Practical *Streptomyces* Genetics. Norwich: John Innes Foundation; 2000.

4. Casu B, Sallmen JW, Schlimpert S, Pilhofer M. Cytoplasmic contractile injection systems mediate cell death in *Streptomyces*. Nat Microbiol. 2023 Apr;8(4):711–26.

5. Gust B, Challis GL, Fowler K, Kieser T, Chater KF. PCR-targeted *Streptomyces* gene replacement identifies a protein domain needed for biosynthesis of the sesquiterpene soil odor geosmin. Proc Natl Acad Sci U S A. 2003 Feb 18;100(4):1541–6.

6. Hong HJ, Hutchings MI, Hill LM, Buttner MJ. The role of the novel Fem protein VanK in vancomycin resistance in *Streptomyces coelicolor.* J Biol Chem. 2005 Apr 1;280(13):13055–61.

7. Schlimpert S, Wasserstrom S, Chandra G, Bibb MJ, Findlay KC, Flärdh K, et al. Two dynamin-like proteins stabilize FtsZ rings during *Streptomyces* sporulation. Proc Natl Acad Sci. 2017 Jul 25;114(30):E6176–83.

8. Karimova G, Robichon C, Ladant D. Characterization of YmgF, a 72-residue inner membrane protein that associates with the *Escherichia coli* cell division machinery. J Bacteriol. 2009 Jan;191(1):333–46.
